# Loci associated with maturation, migration and appetite control are linked with signals of fine-scale local selection in a large Atlantic salmon population

**DOI:** 10.1101/2023.08.23.553800

**Authors:** Antti Miettinen, Johan Dannewitz, Stefan Palm, Ehsan Pashay Ahi, Atso Romakkaniemi, Ville Vähä, Johan Östergren, Craig R. Primmer, Victoria L. Pritchard

**Affiliations:** Organismal & Evolutionary Biology Research Programme, Faculty of Biological and Environmental Sciences, University of Helsinki, Helsinki, Finland; Institute of Biotechnology, University of Helsinki, Helsinki, Finland; Swedish University of Agricultural Sciences, Department of Aquatic Resources, Institute of Freshwater Research, Stångholmsvägen 2, SE-178 93 Drottningholm, Sweden; Natural Resources Institute Finland (Luke), Oulu, Finland; University of the Highlands and Islands, Rivers and Lochs Institute, Inverness, United Kingdom

**Keywords:** Baltic salmon, candidate gene, environmental adaptation, fisheries management, genomic scans, landscape genomics, population genomics, selection signatures

## Abstract

Insights into the genetic basis of local adaptation and drivers of population differentiation improve our understanding of evolution and the maintenance of biological diversity. Characterising adaptively important genetic variation also allows more efficient planning of conservation and management actions. We used a genome-wide SNP array to analyse the genetic population structure of a large Atlantic salmon (*Salmo salar*) population in the interconnected Tornio/Torne and Kalix River system of the Baltic Sea basin, and to identify genomic signatures of fine-scale selection within it. We identified signals of selection and genotype-environment associations (GEA) especially on chromosome (Chr) 9, including on a haploblock containing the *six6* gene and other loci that have been earlier suggested to be adaptively important in salmonids. We also detected signals of selection in genome regions including other genes of ecological relevance, such as two known appetite-controlling genes in the melanocortin system (*pomca* on Chr 9 and *mc4r* on Chr 14), and the maturation-associated gene *taar13c-like* (on Chr 21). Variation in these and other identified candidate genes may potentially reflect differential selective pressures experienced by salmon from different parts of the large river system, regarding traits related to e.g. vision, feeding and growth, age at maturity and/or migratory timing. This indicates a need for management strategies to consider ecologically important genomic regions such as these, in order to protect adaptive genetic diversity in wild salmon populations.

## 1. Introduction

Uncovering the genomic basis of local adaptation is a key aim in evolutionary biology. Adaptive variation determines e.g. the long-term viability of wild populations (e.g. Hohenlohe, Funk, & Rajora, 2021), and the genetic basis of this variation defines how organisms can respond to environmental changes. Thus, understanding how genomic variation contributes to adaptive diversity provides a more complete picture of the evolutionary process and maintenance of biological diversity in general.

Diversity in ecologically relevant traits is important for the adaptive potential (e.g. Hohenlohe et al., 2021) and stability of populations (e.g. Carvalho, Satterthwaite, O’Farrell, Speir, & Palkovacs, 2023; Cordoleani et al., 2021; Gharrett, Joyce, & Smoker, 2013; Hoelzel, Bruford, & Fleischer, 2019), as it can buffer them against environmental change and stochasticity (Schindler et al., 2010). Additionally, it can reduce temporal variability in the numbers of individuals that can be harvested from exploited populations (e.g. Schindler et al., 2010). Characterising adaptively important genetic variation is thus valuable for the conservation and management of threatened natural populations, as knowledge of it can be applied to guide actions that aim to mitigate human-induced disturbances. These applications include identifying population units, predicting their selective responses to anthropogenic effects such as climate change or harvesting regimes, and even management that focuses on specific genetic variants to preserve adaptive diversity of populations (see e.g. Kardos & Shafer, 2018; Shafer et al., 2015; Thompson et al., 2019; Waples et al., 2022).

Population genomics methods allow the identification of genome regions contributing to adaptive variation (e.g. Shafer et al., 2015), ranging from single loci to large structural variants. Signatures of adaptive variation in the genome can be identified by genomic scans that examine spatially varying allele frequencies potentially shaped by selection (reviewed e.g. by Hoban et al., 2016), and by landscape genomic approaches that detect correlations between environmental and genetic variation (e.g. Rellstab, Gugerli, Eckert, Hancock, & Holderegger, 2015). These approaches have been used to identify numerous genomic regions, including large-effect loci associated with important life history variation, as candidates for being under fine- and broad-scale local selection (e.g. Prince et al., 2017; Pritchard et al., 2018).

Atlantic salmon (*Salmo salar*, Salmonidae) is an ecologically, culturally and economically important anadromous fish species that has been heavily impacted by human activities (Myrvold, Mawle, & Aas, 2019; Ignatius, Delaney, & Haapasaari, 2019). Generally, Atlantic salmon spend their first years in flowing fresh water, from where they migrate to the sea, returning one to multiple years later to spawn near their natal freshwater location (Thorstad et al., 2011; Webb et al., 2007). This homing behaviour can promote reproductive isolation (e.g. Quinn, 1993), thus providing opportunities for locally adaptive differentiation over fine spatial scales. Fine-scale, putatively adaptive genetic variation has indeed been identified in recent studies of different Atlantic salmon lineages (e.g. Pritchard et al., 2018; Wellband et al., 2018; Watson et al., 2022).

A distinct phylogeographic lineage of Atlantic salmon migrates into the Baltic Sea, one of the world’s largest bodies of brackish water, located in northern Europe. In the past century, hydropower dam construction, pollution and overfishing have caused the loss of Baltic salmon populations from approximately two thirds of the c. 100 rivers where they historically spawned in (Karlsson & Karlström, 1994; Palmé, Wenneström, Guban, Ryman, & Laikre, 2012; Romakkaniemi et al., 2003). Stocking of hatchery-origin salmon sustains many of the remaining stocks (ICES, 2023). Increased gene flow due to stocking has resulted in more homogeneous salmon populations (both within and among rivers) over the last century, raising concern for the ability of the remaining populations to adapt to changing environmental conditions (Östergren et al., 2021). Many of the remaining wild populations in the northernmost part of the Baltic Sea (the Gulf of Bothnia) have recovered since the late 1990s, and salmon from these rivers are actively harvested both at sea and in rivers (ICES, 2020). The marine fishery represents one of the few remaining commercial sea harvests of Atlantic salmon currently permitted in the world. To safeguard the adaptive potential of these salmon stocks, their adaptive genetic diversity needs to be examined.

The largest remaining wild Baltic salmon stock spawns in the unregulated Tornio (Torne in Swedish) and Kalix River complex (ICES, 2023); two large neighbouring river systems partly interconnected by a bifurcation (Figure 1). Because of their long migrations passing through all the major salmon fishing areas in the Baltic Sea, salmon from these rivers are heavily exploited by fisheries (ICES, 2023). An earlier study using 18 microsatellite markers found that the population structure of salmon in this large river system was linked to life history variation: the main genetic divergence in the system was between salmon from upper and lower river sections, and it was associated with differences in smolt and adult migration timing (Miettinen et al., 2021). However, neither the factors potentially driving population structuring nor possible adaptive differentiation in this system have been previously investigated.

**Figure 1.**
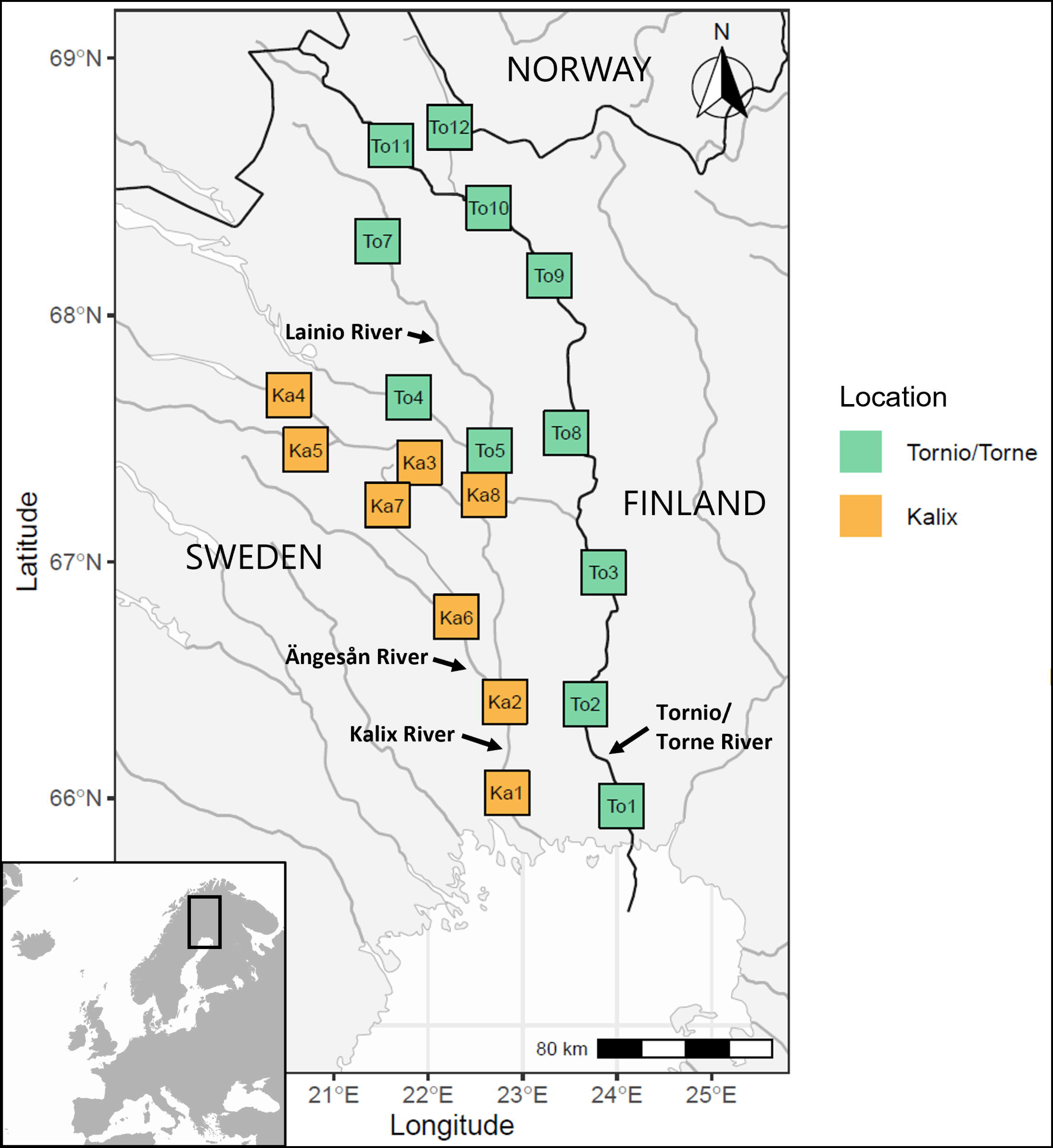
Map showing the location of samples used in the study, collected from 19 different sites in the Tornio (in Finland/Sweden) and Kalix (in Sweden) Rivers. A bifurcation (Tärendö River) connecting the Tornio and Kalix is located at site Ka8. The darker line depicts national borders, including the Tornio River that flows on the border of Finland and Sweden.

In this study, we used a genome-wide dataset from a SNP array to assess the fine-scale genetic structuring of Atlantic salmon in the Tornio-Kalix system and search for genomic signatures of local selection, possibly associated with environmental variation. The aim was to improve our understanding of adaptive genetic diversity and its potential drivers in this important wild salmon stock, and generate information that may help to understand the possible responses of this stock to human-induced selective pressures and environmental changes.

## 2. Material and methods

### 2.1 Study area and sample collection

The Tornio-Kalix River complex contains the Tornio (Torne in Swedish) River that is one of the largest unregulated rivers in northern Europe (length 522 km, watershed area 40,157 km2, mean discharge 389 m^3s-1^, c. 50,000-150,000 returning spawners annually in recent years), and the Kalix River (length 461 km, watershed area 23,600 km2, mean discharge 299 m ^3s-1^, c. 30,000-60,000 returning spawners annually in recent years), that are located in a terrain that ranges from a boreal zone to a subarctic headwater zone (HELCOM, 2011; Romakkaniemi et al., 2003; ICES, 2023). The two rivers have their mouths located c. 50 km apart, and flow into the northernmost part of the Gulf of Bothnia in the Baltic Sea. A natural bifurcation (Tärendö River) ca. 200-250 km upstream of the river mouths connects the two rivers, allowing more than half of the annual discharge of the Tornio (Torne) River main branch to flow into the Kalix River main stem (e.g. Dankers & Middelkoop, 2008).

Tissue samples of wild salmon juveniles were collected from 52 electrofishing locations across the Tornio (n = 221, years 2012-2014) and Kalix (n = 155, years 2012-2015) Rivers (Table 1). The samples used here (n = 376) consisted of a subset (n = 282) of the juveniles used in Miettinen et al. (2021), 61 parr from three new sites in the Tornio River (sites To9, To10 and To11) and 34 parr from two sites in the Kalix River (sites Ka6 and Ka7) (Table S1). We merged the locations into 19 broader sampling sites on the basis of geographical proximity (Figure 1, Table 1), as in Miettinen et al. (2021). The number of sampled juveniles per site ranged from 17 to 21.

**Table 1.** Sampling site information.

| Sampling site | Location | Latitude (°N) | Longitude (°W) | Samples (n)* | Sampling years |
| --- | --- | --- | --- | --- | --- |
| **Ka1** | Forsbyn | 66.0758679 | 22.7313647 | 8 | 2012 |
|  | Forshaga | 65.9501012 | 22.874327 | 6 |  |
|  | Kamlunge | 66.0025135 | 22.8499411 | 5 |  |
|  | Rian | 65.8809964 | 23.0048283 | 1 |  |
| **Ka2** | Ansvarforsen | 66.613788 | 22.7522401 | 11 | 2012 |
|  | Orrforsen | 66.5562608 | 22.7508014 | 10 |  |
| **Ka3** | Mustisuantto | 67.3692504 | 21.9365067 | 8 | 2012 |
|  | Nurmikoski | 67.3921581 | 21.8928112 | 8 |  |
|  | Nurmisuantto | 67.4059321 | 21.9117864 | 4 |  |
| **Ka4** | Kalixfors | 67.7337232 | 20.323975 | 10 | 2012 |
|  | Saarisuantto | 67.6752691 | 20.5557125 | 9 |  |
| **Ka5** | Akkalkoski | 67.471751 | 20.5688603 | 8 | 2012 |
|  | Laxselet | 67.4773639 | 20.5369344 | 5 |  |
|  | Luspakoski | 67.4455858 | 20.8757275 | 7 |  |
| **Ka6** | Kurkkiokoski | 66.8982004 | 22.0041706 | 16 | 2015 |
| **Ka7** | Niilivaarabron | 67.1917784 | 21.6024237 | 17 |  |
| **Ka8** | Pistoniva | 67.343954 | 22.4410378 | 11 | 2012 |
|  | Pitkäkoski | 67.1884331 | 22.7298243 | 9 |  |
| **To1** | Kiviranta | 65.8516998 | 24.1629342 | 4 | 2012 |
|  | Oravaisensaari | 65.9185498 | 24.1270009 | 6 |  |
|  | Tanskinsaari | 65.8820998 | 24.1476509 | 5 |  |
|  | Vähänärä | 65.9325831 | 24.0980843 | 4 |  |
| **To2** | Korpikoski | 66.7017276 | 23.8994462 | 4 | 2012-2013 |
|  | Matkakoski | 66.142041 | 23.9334641 | 5 |  |
|  | Turtola | 66.6281485 | 23.8783968 | 4 |  |
|  | Vuennonkoski | 66.1700562 | 23.8466294 | 4 |  |
| **To3** | Jarhoinen | 66.952267 | 23.8643357 | 6 | 2012 |
|  | Kaartisenniva | 67.4479508 | 23.5122696 | 7 |  |
|  | Kassa | 67.0681505 | 23.6736191 | 7 |  |
| **To4** | Jölketturkkio | 67.690781 | 21.4304608 | 1 | 2012 |
|  | Saarikonsuantto | 67.7756797 | 20.9265822 | 7 |  |
|  | Talvimaansuantto | 67.736168 | 21.0184749 | 7 |  |
|  | Tervaskoski | 67.7346393 | 21.1679981 | 5 |  |
| **To5** | Kangosenkoski | 67.4478073 | 22.6617762 | 10 | 2012 |
|  | Sienikoski | 67.5093604 | 22.6632245 | 10 |  |
| **To7** | Kenttäkoski övre | 68.2920307 | 21.451032 | 10 | 2012 |
|  | Suunsaari yttre | 68.2785338 | 21.4436546 | 10 |  |
| **To8** | Kaarnekoski | 67.6272176 | 23.5412366 | 8 | 2012 |
|  | Kihlanki | 67.5641842 | 23.4881197 | 4 |  |
|  | Mukkaskoski | 67.4479508 | 23.5122696 | 3 |  |
|  | Pyssykorva | 67.6883509 | 23.49302 | 4 |  |
| **To9** | Sonkamuoika | 68.148883 | 23.244183 | 14 | 2012-2014 |
|  | Visanto | 68.02955 | 23.467933 | 4 |  |
| **To10** | Jatuni | 68.431667 | 22.645083 | 15 | 2012-2014 |
|  | Kuttanen | 68.380333 | 22.846267 | 4 |  |
| **To11** | Lammaskoski | 68.766117 | 21.378117 | 4 | 2012-2014 |
|  | Naimakkaluspa | 68.661333 | 21.624533 | 6 |  |
|  | Vuokkasenniva | 68.5853 | 21.870617 | 7 |  |
| **To12** | Kinnerpuska | 68.7491356 | 22.209137 | 5 | 2012 |
|  | Mukkakoski | 68.7687356 | 22.1881036 | 6 |  |
|  | Patoniva | 68.7326023 | 22.2345703 | 1 |  |
|  | Pultsakurkkio | 68.7358189 | 22.224487 | 7 |  |

\* Individuals used in analyses.

### 2.2 Environmental, geographic and bioclimatic data

We collected information of 26 environmental, geographic and bioclimatic variables available for the sampling locations in our dataset (Table S2). For each variable, we used the weighted average of measurements from each location for each site (based on the number of samples per location). We then standardised the variables by subtracting the mean and dividing by the standard deviation, using the *scale()* function in R 4.1.3 (R Core Team, 2022). We tested for correlations among these variables using a Pearson correlation test in R and retained only 10 variables with a Pearson correlation coefficient (r) of < 0.7 with any of the other (nine) retained variables. The retained variables were distance from river mouth, water depth, water velocity, bottom substrate, underwater vegetation, isothermality, temperature annual range, mean temperature of wettest quarter, precipitation of the wettest quarter, annual precipitation (Figure S1). We note that distance from river mouth was very strongly correlated with elevation, latitude and mean annual temperature (not retained; Figure S2) and can therefore be considered a temperature-related proxy in our analyses.

### 2.3 DNA extractions

We extracted DNA using the QIAGEN DNAamp Mini Kit method, the QuickExtract™ DNA Extraction Solution method (Lucigen), or a salt extraction method (Aljanabi & Martinez, 1997). We used a NanoDrop ND-1000 spectrophotometer (Thermo Fisher Scientific) and Qubit fluorometer with the dsDNA BR Assay Kit (Thermo Fisher Scientific) to assess DNA concentrations. We then standardised the DNA to a NanoDrop-estimated concentration of 25-35 ng/µl.

### 2.4 Genotyping and data quality filtering

Genotyping was conducted at the Centre for Integrative Genetics (CIGENE, Norwegian University of Life Sciences, Norway) with a custom SNP microarray (Affymetrix Axiom) containing 60,252 markers developed for Atlantic salmon. Samples with a dish quality control (DQC) metric < 0.95 and a call rate < 0.97 were removed from downstream analyses (n = 7). To assess and control for sequencing batch effects among genotyping runs, we genotyped the same individuals (n = 13) on all runs.

First, we mapped the positions of the SNP markers to the Ssal_v3.1 reference genome (assembly GCA_905237065.2). Overall, 54,763 SNPs mapped to the reference, with the total genotyping rate being 98.3%. Next, using PLINK v1.9 (Chang et al., 2015) we filtered out SNPs that did not map to chromosomes 1-29 on the reference genome (n = 642). We then filtered out SNPs with i) any mismatching genotypes in control samples among runs (n = 2,772); ii) > 10% missing data (n = 1,262); iii) a minor allele frequency (MAF) < 5% (n = 12,768); and iv) strong deviations from Hardy-Weinberg equilibrium (HWE p < 0.00001; n = 64), possibly indicative of technical genotyping problems. We based the HWE calculation only on individuals from the lower parts of the Tornio-Kalix system (n = 174; excluding the bifurcation site Ka8) that comprised the majority of samples in the dataset and were known to not contain significant substructure (Miettinen et al., 2021). This filtered dataset used for downstream analyses consisted of 37,255 SNPs and had an overall genotyping rate of 99.2%.

To avoid family sampling of juvenile salmon that can bias allele frequency estimates (Hansen, Nielsen, & Mensberg, 1997; Hansen & Jensen, 2005; Östergren, Palm, Gilbey, & Dannewitz, 2020), we studied the presence of closely related individuals in the dataset by splitting it into the different sampling sites, applying an additional MAF filter (< 0.05) for each site, and using the *--genome* function in PLINK to estimate pairwise genome-wide identity-by-descent (IBS) between each individual. We then used a PI_HAT (pairwise IBS value) threshold of > 0.35 to identify potential close relatives, and retained only one individual per group of putative full siblings. In total we removed seven individuals from putative full-sib pairs.

We used PLINK or PGDSPIDER version 2.1.1.5 (Lischer & Excoffier, 2012) to convert input files into different formats when needed. We used BEAGLE 5.4 (Browning, Tian, Zhou, & Browning, 2021; Browning, Zhou, & Browning, 2018) with the filtered dataset to impute missing genotypes or infer phasing where necessary for our analyses. Two SNPs were found to share the same chromosomal position, so we removed one of them from the phased dataset.

### 2.5 Assessing genetic diversity and population structure

#### 2.5.1 Expected and observed heterozygosity

We used *adegenet* 2.1.10 (Jombart & Ahmed, 2011) and *hierfstat* 0.5-11 (Goudet & Jombart, 2022) in R to estimate genome-wide expected and observed heterozygosities for each sampling site, over all filtered SNPs (n = 37,255).

#### 2.5.2 Runs of homozygosity

We used PLINK to analyse runs of homozygosity (ROH) in the dataset, to compare among-site variation in the length distributions of ROH, which can be informative about population history (long ROH may indicate isolated or bottlenecked populations) (e.g. Ceballos, Joshi, Clark, Ramsay, & Wilson, 2018; Foote et al., 2021; Purfield, Berry, McParland, & Bradley, 2012). We used the following settings in PLINK: *-- homozyg --homozyg-density 50 --homozyg-gap 500 --homozyg-window-snp 50 --homozyg-window-threshold 0.5*. We used the mean length of ROH in samples from each sampling site.

#### 2.5.3 Population structure

For studying the genetic population structure in the Tornio-Kalix system, we removed SNPs in high linkage disequilibrium (n = 16,510) using PLINK (command *--indep-pairwise 50 5 0.5*). This “LD-pruned dataset” for analyses of population structure consisted of 20,745 SNPs, with a total genotyping rate of 99.2%. We used this LD-pruned dataset i) to estimate the number of distinct genetic clusters by running ADMIXTURE 1.3.0 (Alexander, Novembre, & Lange, 2009) with *K* = 1 to *K* = 19, using the five-fold cross-validation (CV) error for each *K* to determine the optimal number of clusters, and ii) to perform a principal component analysis (PCA) with PCADAPT 4.3.2 (Privé, Luu, Vilhjálmsson, Blum, & Rosenberg, 2020) in order to determine the optimal number of principal components (PCs) to describe the data. We visualised the ADMIXTURE results with POPHELPER (Francis, 2017) in R, and assessed the optimal number of PCs by visually inspecting screeplots and PCA plots using *ggplot2* (Wickham 2016) in R (Figure S4b). Furthermore, we used iii) *StAMPP* (Pembleton, Cogan, & Forster, 2013) in R to estimate global and pairwise F_ST_.

### 2.6 Identifying genomic signatures of local selection

As a combination of different approaches is recommended for identifying candidate genome regions under putative selection (e.g. de Villemereuil, Frichot, Bazin, François, & Gaggiotti, 2014; Rellstab et al., 2015), we applied several analytical frameworks (described in Table 2) on the dataset of 37,255 SNPs, and only discuss regions showing signals of local selection in multiple types of analyses.

**Table 2.** Description of methods and their test statistics used in this study to detect signals of local selection.

| Approach | Software | Test statistic | Method description | Definition of outliers* |
| --- | --- | --- | --- | --- |
| Genome scans for outliers | *pcadapt 4.3.2*<br>(Privé et al., 2020) | Mahalanobis distances, p-value | Identifies outliers based on correlations between each genetic marker and population structure determined by the optimal number of principal components. | “Outlier-based candidates” were SNPs in the top-ranked 0.5% in all of the tests. |
|  | *BayPass 2.31*<br>(Gautier, 2015) | Absolute XtX | Bayesian method that identifies outliers by estimating per-marker deviations from the underlying genome-wide population structure by using a neutral covariance matrix of population allele frequencies. |  |
|  | *BayeScEnv v1.1*<br>(de Villemereuil & Gaggiotti, 2015) | q-value, alpha parameter | Bayesian method that estimates the posterior probability that a given locus is under selection. |  |
| Univariate genotype-environment association (GEA) | *LFMM 2* (Caye, Jumentier, Lepeule, & François, 2019) | q-value | Detects associations of allele frequencies with environmental variables while using latent factors to account for population structure. | “Univariate GEA candidates” for each environmental variable were SNPs in the top-ranked 0.5% in all of the tests. |
|  | *BayPass 2.31* | Absolute Pearson correlation coefficient (r) | Bayesian method that identifies loci outliers associated with population-specific covariates, while accounting for underlying genome-wide population structure. |  |
| | *BayeScEnv v1.1* | g, q-value | Bayesian method that identifies outlier loci showing an increase in between-population $F_{ST}$ with increasing environmental differentiation. | |
| Multivariate GEA (redundancy analysis; RDA) | *vegan 2.6-4*<br>(Oksanen et al., 2022) | Loadings in the ordination space | PCA-based method that determines how sets of loci covary in response to the multivariate environment, and can detect relatively weak, multilocus signatures of selection (Forester et al., 2018). Partial RDA can be used to correct for population structure. | “pRDA/full RDA candidates” had extreme loadings on the RDA axes. |
| Haplotype-based | *selscan v1.3.0*<br>(Szpiech & Hernandez, 2014) | XP-EHH (maximum absolute normalised score) | Identifies genomic regions suggestive of hard selective sweeps, indicated by elevated haplotype homozygosity. | “XP-EHH candidates” were SNPs the top-ranked 0.5%. |

\* In this study.

Following Pritchard et al. (2018) and Zueva, Lumme, Veselov, Primmer, & Pritchard (2021), we ranked SNPs by test score and considered the 0.5% highest-ranked SNPs (i.e. 186 SNPs; equivalent to an empirical p-value < 0.005 when considering all SNPs in the dataset) from each test as putative outliers in all analyses except for the multivariate GEA.

#### 2.6.1 Outlier-based approaches

We used three outlier-based approaches to identify highly differentiated loci among sampling sites, consistent with locally divergent selection (Table 2). We used i) PCADAPT in R with three principal components, based on our population structure analysis (2.5.3 above), ii) BAYPASS to first generate five independent covariance matrices using the LD-pruned SNP dataset (n = 20,745; non-default parameters used were *-npilot 20 -pilotlength 1000 -nthreads 8*) and then conduct five replicate runs (non-default parameters used: *-npop 19 -npilot 25 -pilotlength 2000 -nval 2000 -nthreads 8*) using a different covariance matrix to control for population structure for each run, and iii) B AYESCENV with a model not including environmental associations (equivalent to a BayeScan model, Foll & Gaggiotti, 2008) (non-default parameters used: *-threads 16 -pr_pref 1 -pr_jump 0.005 -nbp 12*). All the BAYPASS runs were initialised with a different random number seed, and we used the median XtX value over the five runs for each SNP as their test statistic. For B AYESCENV that attempts to infer both divergent and balancing selection, we only considered SNPs with a positive alpha parameter value, indicative of divergent selection.

#### 2.6.2 Genotype-environment association (GEA) analyses

##### 2.6.2.1 Univariate GEA

We used three genotype-environment association (GEA) approaches that perform univariate statistical analyses to identify correlations between genetic variation and the 10 environmental variables (Table 2): i) BAYPASS as above (2.6.1), except for using the median Pearson correlation coefficient (r) over the five runs for each SNP as their test statistic, ii) B AYESCENV as above (2.6.1), except for using the parameter *-pr_pref 0*, and iii) LFMM using the function *lfmm_ridge* and *K* = 4 latent factors, based on the number of genetic clusters identified when analysing population structure (2.5.3 above).

##### 2.6.2.2 Multivariate GEA

We used redundancy analyses (RDA) with the R package *vegan* to determine how sets of loci covary in response to the multivariate environment (i.e. the 10 environmental variables we used). To control for population structure that may cause spurious detections of candidate loci caused by neutral processes (e.g. de Villemereuil et al., 2014; Excoffier, Hofer, & Foll, 2009), we performed a partial RDA (pRDA) using values of the first three PC axes that best explained population structure (see 2.5.3) as a conditioning matrix. Because correcting for population structure can also remove true selective signals of environmental variables that co-vary with the structure (Forester, Lasky, Wagner, & Urban, 2018), we also performed a full RDA not accounting for it. Based on inspection of screeplots and distribution of p-values (as in Blanco-Pastor et al., 2021), we retained the first two RDA axes in both pRDA and full RDA. We identified candidate SNPs (“pRDA candidates” from the pRDA and “full RDA candidates” from the full RDA) by using a relatively liberal cut-off of loadings > 3 SD from the mean distribution of each of the two RDA axes (as in e.g. Forester et al., 2018; Salisbury et al., 2023) in each RDA, corresponding to a two-tailed p-value threshold of 0.0027.

Furthermore, we used variance partitioning in partial RDA to gain estimates of the contributions of defined sets of variables on the total observed genetic variation in the dataset (Table S4), i.e. to study the independent impacts of geography, environment, and past demographic processes (neutral population structure) on the observed genetic variation when the influence of other variables has been removed (reviewed in Capblancq & Forester, 2021; Forester et al., 2018).

#### 2.6.3 Haplotype homozygosity

We investigated cross-population extended haplotype homozygosity (XP-EHH) (Sabeti et al., 2007) at each SNP to identify genomic regions suggestive of hard selective sweeps, indicated by elevated haplotype homozygosity. Based on our analysis of population structure (see 2.5.3), we classified each sampling site to one of the identified genetic clusters (subpopulations), and then used SELSCAN with default settings (except for *--max-gap 2000000*, as in Pritchard et al., 2018) to estimate XP-EHH for each SNP for each of the six pairwise subpopulation comparisons. Then, we standard-normalised the XP-EHH values across all chromosomes and within each pairwise comparison (using function *--norm*), and used the maximum absolute normalised score over all comparisons as our test statistic for each SNP (as in Pritchard et al., 2018).

#### 2.6.4 Identification of candidate haploblocks

We used PLINK (function *--blocks*) to identify sets of neighbouring SNPs in high linkage disequilibrium (hereafter “haploblocks” or “blocks”), in order to assess if multiple of our candidate SNPs could be tagging the same locus. Following Pritchard et al. (2018), we used the following parameters for haploblock identification: *--no-small-max-span --blocks-inform-frac 0.8 --blocks-max-kb 5000 --blocks-strong-lowci 0.55 --blocks-strong-highci 0.85 --blocks-recomb-highci 0.8*. We then defined the haploblock boundaries as the genomic positions halfway between the blocks’ outermost SNPs and the closest SNPs outside of it. Any haploblocks containing candidate SNPs closer than 10kb apart were condensed into a single block. To avoid false positives, we used a conservative approach of classifying the detected blocks as “candidate haploblocks” and discussing them only if they contained candidate SNPs from multiple types of analyses (i.e. candidates from at least two of the following: outlier-based analyses, GEA analyses, XP-EHH testing).

#### 2.6.5 Annotation of candidate genomic regions

We used information from the NCBI Salmo salar Annotation Release 102, and the *intersect* function of BEDTOOLS 2.30.0 (Quinlan & Hall, 2010) to annotate genes within the candidate haploblocks. Then, we made a literature search to assess whether these genes have been found relevant for local adaptation to environmental conditions in other studies of salmonids and other fishes.

#### 2.6.6 Assessing overlap with previously identified structural variants

Structural variants are increasingly recognized as being adaptively important in many taxa (e.g. Cayuela et al., 2020; Fuentes-Pardo, Farrell, Pettersson, Sprehn, & Andersson, 2023; Han et al., 2020; Le Moan, Bekkevold, & Hemmer-Hansen, 2021), and have been shown to be common in Atlantic salmon genomes (Bertolotti et al., 2020). We checked our candidate genomic regions for overlap with previously identified structural variants in Atlantic salmon (Bertolotti et al., 2020) with BEDTOOLS, using the function *intersect*.

## 3. Results

### 3.1 Genotyping and data quality filtering

In total, 369 individuals ranging from 16 to 21 per sampling site (Table 1) met the Affymetrix sequencing quality thresholds. We removed seven individuals from full-sib pairs, and one individual (from site Ka4) that was suspected to be a sample mix-up not originating from the river system based on initial screenings of population structure (not shown). The final dataset for downstream analyses consisted of 361 individuals.

### 3.2 Genetic diversity and population structure

Observed heterozygosity (Hobs) per sampling site ranged from 0.337 (Ka4) to 0.38 (To2). In general, sites in the main stems of the lower Tornio and Kalix showed higher Hobs than those in the Ängesån tributary or upper parts of the river system (Table S1). Runs of homozygosity (ROH) were on average longer for individuals sampled from the Ängesån tributary and upper parts of the system, indicating higher levels of inbreeding (Table S1).

ADMIXTURE results (cross-validation error, CV) for the LD-pruned dataset suggested that the optimal *K* number of clusters was 3, but CV errors of *K* values from 2 to 4 were close (Figure S3, Figure S4a). *K* = 4 was best in line with geography, as four genetic clusters could distinguish the lower and upper parts of the river system, as well as two tributary populations (the upper parts of the Lainio River in the Tornio/Torne River system, and the Ängesån River in the Kalix River system) (Figure 2). The same four genetic units were supported by PCA that showed three PCs to explain most of the variation in the system (Figure 2, Figure S4b). The main genetic cluster consisting of most downstream and upstream individuals followed geography quite closely, with the geographically most distant sites being far apart on the PCA plot. A small number of individuals collected from the lower parts of the river system genetically resembled individuals from upper parts of the system.

**Figure 2.**
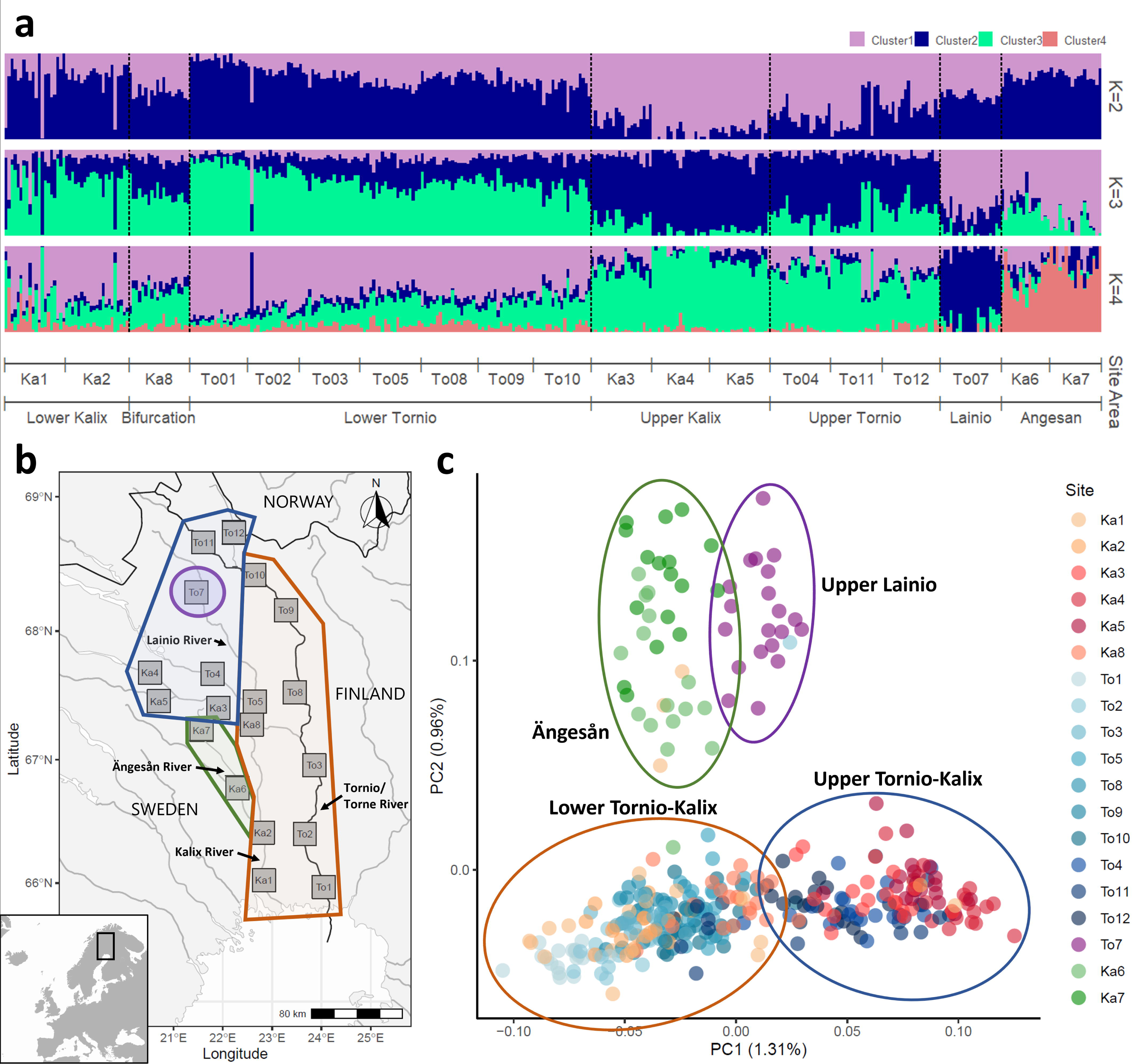
**a** ADMIXTURE coefficients for the Tornio-Kalix salmon at*K* values from 2 to 4 (each vertical line depicts an individual), **b** a map of the Tornio-Kalix River system, with the identified genetic clusters of the salmon stock overlaid on it, and **c** a principal component analysis (PCA) visualising the genetic population structure of the Tornio-Kalix salmon stock, showing the first two PC axes. The proportion of variance explained by each principal component is indicated on the axis labels. Each point represents an individual, and their distribution on the axes depicts their genetic distance from each other. ADMIXTURE and PCA were run using the LD-pruned dataset of 20,745 SNPs.

Global F _ST_ across all sites was 0.015. Pairwise F _ST_ ranged from 0.001 (between Ka1 and Ka2) to 0.042 (between Ka4 and Ka7). The pairwise F _ST_ estimates between sites in the upper Lainio or Ängesån tributary versus any other sites ranged from 0.022 (between To7 and Ka8) to 0.039 (between To7 and Ka4), and from 0.011 (between Ka6 and Ka1) to 0.042 (between Ka7 and Ka4) (Table S3).

### 3.3 Genomic signatures of local selection

#### 3.3.1 Outlier-based analyses

Out of the top-ranked SNPs (0.5% of the dataset, i.e. 186 SNPs), 90, 82 and 98 were found on Chromosome 9 using PCADAPT, BAYPASS, and BAYESCENV, respectively (48.4%, 44.1% and 52.7% of the top-ranked outliers in these analyses, respectively). Overall, 65 SNPs were top-ranked in all three of the outlier-based analyses (hereafter referred to as “outlier-based candidates”, Figure S5), of which 59 (90.8%) were on Chromosome 9, one on Chr 13, one on Chr 14, two on Chr 20 and one on Chr 28.

#### 3.3.2 Genotype-environment association (GEA) analyses

##### 3.3.2.1 Univariate GEA analyses

We identified 84 SNPs that were top-ranked across all three different univariate GEA analyses for each variable (hereafter “univariate GEA candidates”). Of those, 35 were associated with bottom substrate, 27 with BIO3, 23 with distance from river mouth, nine with annual precipitation, eight with BIO16, seven with vegetation, five with BIO8, six with depth, and four with velocity. Twenty-eight of these “univariate GEA candidates” were associated with multiple environmental variables. The highest number of univariate GEA candidates were on Chr 9 (13 out of the 84 SNPs, i.e. 15.5%).

##### 3.3.2.2 Multivariate GEA analysis (RDA)

Using a partial redundancy analysis (pRDA) to control for population structure, and a “full” RDA not accounting for it, we identified 371 SNPs (hereafter “pRDA candidates”) and 432 SNPs (hereafter “full RDA candidates”) associated with multivariate environmental variation, respectively. Only six of these 803 candidate SNPs were shared among the two different types of RDA (Figure S5). Chr 9 contained the highest number of candidates in the full RDA (131 out of 432 SNPs, i.e. 30.3%) and second most (after Chr 1) in the pRDA (27 out of 371 SNPs, i.e. 7.3%). The most correlated predictors of the pRDA candidates and full RDA candidates, respectively, were as follows: bottom substrate (n = 91, n = 32), distance from river mouth (n = 54, n = 71), water velocity (n = 37, n = 52), BIO16 (n = 30, n = 175), BIO3 (n = 29, n = 74), underwater vegetation (n = 28, n = 1), BIO8 (n = 27, n = 3), water depth (n = 26, n = 32), BIO12 (n = 26, n = 14), and BIO7 (n = 23, n = 3).

Partitioning the observed genetic variance into different components showed that geography/environment, climate and genetic structure together explained 6.9% of the overall genetic variance across the river system, while the fraction of unexplained variation was 93.1% (Table S4). Geography/environment and climate independently explained 1.7% and 1.8% of the total genetic variation, respectively (i.e. 25.4% and 24.7% of the variance explained by the full RDA model, respectively) when population structure was controlled for (Table S4). The genetic structure alone explained 1.7% of the total variance (i.e. 25.2% of the total explained variation). The confounded effect of the different components (i.e. geography/environment, climate and genetic structure) also accounted for a fourth of the total explained variation (24.7%), meaning that this portion of genetic variance could not be uniquely associated with any of the three sets of predictors.

#### 3.3.3 Haplotype homozygosity

The haplotype-based XP-EHH analysis showed that the 186 top-ranked SNPs showing signs of elevated haplotype homozygosity (hereafter “XP-EHH candidates”), possibly indicative of a selective sweep on an allele within the haplotype, were mostly found on Chr 9 (112/186, 60.2% of all candidates). Out of the XP-EHH candidates, 20 overlapped with the 65 outlier-based candidates (19 of them on Chr 9, one on Chr 28), two with the 84 univariate GEA candidates, and 30 with the 803 RDA candidates (Figure S5).

#### 3.3.4 Candidate haploblocks

We found 311 unique candidate SNPs from outlier-based analyses, univariate GEA analyses, and XP-EHH testing (Figure S5), contributing to 105 unique haploblocks that contained these candidate SNPs. The length of the 105 blocks ranged from 0.004 Mb to 4.45 Mb (shortest and longest haploblocks were both on Chr 5).

The RDA analyses identified 693 additional unique candidate SNPs. The 105 haploblocks mentioned above contained 159 RDA candidate SNPs (44 from pRDA, 109 from f2RDA, 6 from both), with 103 of them matching SNPs identified by other analyses (Figure S5).

Focusing only on candidate genomic regions with strongest support for being putatively under selection, we discuss only haploblocks containing SNPs identified by multiple types of analyses (i.e. at least two of the following: outlier-based, GEA or XP-EHH candidates; therefore ignoring blocks containing only univariate GEA and RDA candidates). We found 18 such haploblocks (hereafter “candidate haploblocks”) (Table 3; Figures S6a-r). The highest number (n = 8) of these blocks was on Chr 9. The environmental variables associated with these candidate blocks in either univariate or multivariate GEA analyses were BIO16 (nine times), distance from river mouth (six times), BIO3 (six times), depth (three times), velocity (twice), BIO7 (once), BIO12 (once), bottom substrate (once), and underwater vegetation (once). Three of the blocks included candidate SNPs from all different types of analyses (i.e. from outlier-based, univariate and multivariate GEA, and XP-EHH tests), all of them on Chr 9. The size of the candidate blocks ranged from 0.05 Mb (on Chr 28) to 2.37 Mb (on Chr 18). Out of the 223 genes found within these candidate blocks, 161 were on Chr 9 (Table 3). Overall, these 18 blocks contained 164 of the 311 candidate SNPs identified by the outlier-based, univariate GEA or XP-EHH approaches, and 104 of the 803 RDA candidates. We list putatively adaptive genes within the candidate haploblocks in Table S5. Figure 3 shows empirical p-values of test scores, excluding scores based on RDA loadings, and annotated genes in the haploblock containing most candidate genes (block 2), and Figures S5a-r show these for all the 18 candidate haploblocks.

**Table 3.** Information about the 18 candidate haploblocks identified in this study. Type refers to the approach that identified candidates in each block (“GEA (both)” = identified by both univariate and multivariate GEA; “RDA (both)” = identified by both pRDA and full RDA), and GEA refers to the environmental variable associated with each block (associations identified by both univariate and multivariate GEA are underlined).

| Block # | Chr | Start (bp) | End (bp) | Length (Mb) | Type | GEA<sup>a</sup> | # candidate SNPs (and genes) | Structural variants<sup>b</sup> |
| --- | --- | --- | --- | --- | --- | --- | --- | --- |
| 1 | 9 | 22041594 | 22229310 | 0.19 | RDA, XP-EHH | Distance | 3 (6) |  |
| 2 | 9 | 22646469 | 24333545 | 1.69 | Outlier, GEA (both), XP-EHH | <u>BIO12</u>, BIO16, distance, velocity | 26 (50) |  |
| 3 | 9 | 24394915 | 25479569 | 1.08 | Outlier, RDA (both), XP-EHH | BIO3, BIO16, depth, distance, velocity | 59 (29) |  |
| 4 | 9 | 25544576 | 26572371 | 1.03 | Outlier, GEA (both), XP-EHH | Depth, <u>distance</u> | 41 (29) | One candidate SNP in a 860bp deletion (Bertolotti et al., 2020). |
| 5 | 9 | 28118150 | 28415589 | 0.30 | Outlier, RDA (both), XP-EHH | BIO16 | 5 (1) |  |
| 6 | 9 | 28487961 | 28693554 | 0.21 | Outlier, GEA (both), XP-EHH | <u>BIO3</u>, BIO16 | 4 (6) |  |
| 7 | 9 | 28708901 | 28998334 | 0.29 | Outlier, GEA (both) | <u>BIO16</u> | 4 (7) |  |
| 8 | 9 | 44000579 | 46193652 | 2.19 | Outlier, RDA | BIO3, BIO16 | 10 (33) | One candidate SNP in a 3110bp deletion (Bertolotti et al., 2020). |
| 9 | 13 | 33821031 | 34113068 | 0.29 | Outlier, RDA | BIO16 | 1 (4) |  |
| 10 | 13 | 72834066 | 73244495 | 0.41 | RDA, XP-EHH | BIO3 | 6 (7) |  |
| 11 | 14 | 24272662 | 24610951 | 0.34 | Outlier, RDA | BIO16 | 1 (9) |  |
| 12 | 14 | 31661331 | 31822502 | 0.16 | RDA (both), XP-EHH | BIO7, distance | 3 (6) |  |
| 13 | 14 | 47642849 | 48234437 | 0.59 | RDA, XP-EHH | BIO3 | 4 (8) |  |
| 14 | 17 | 56717526 | 56866325 | 0.15 | RDA, XP-EHH | Distance | 3 (6) |  |
| 15 | 18 | 1235854 | 3603311 | 2.37 | GEA (both), XP-EHH | <u>Bottom substrate, vegetation</u> | 4 (9) |  |
| 16 | 20 | 27662780 | 27760237 | 0.10 | Outlier, RDA | BIO3 | 2 (5) |  |
| 17 | 21 | 49415626 | 49962816 | 0.55 | Outlier, GEA (both) | <u>BIO16</u> | 3 (6) |  |
| 18 | 28 | 6645841 | 6691045 | 0.05 | Outlier, RDA, XP-EHH | Depth | 2 (2) |  |

<sup>a</sup> BIO3 = Isothermality, BIO7 = Temperature annual range, BIO12 = Annual precipitation, BIO16 = Precipitation of wettest quarter.
<sup>b</sup> Structural variants that directly overlapped with candidate SNPs.

**Figure 3.**
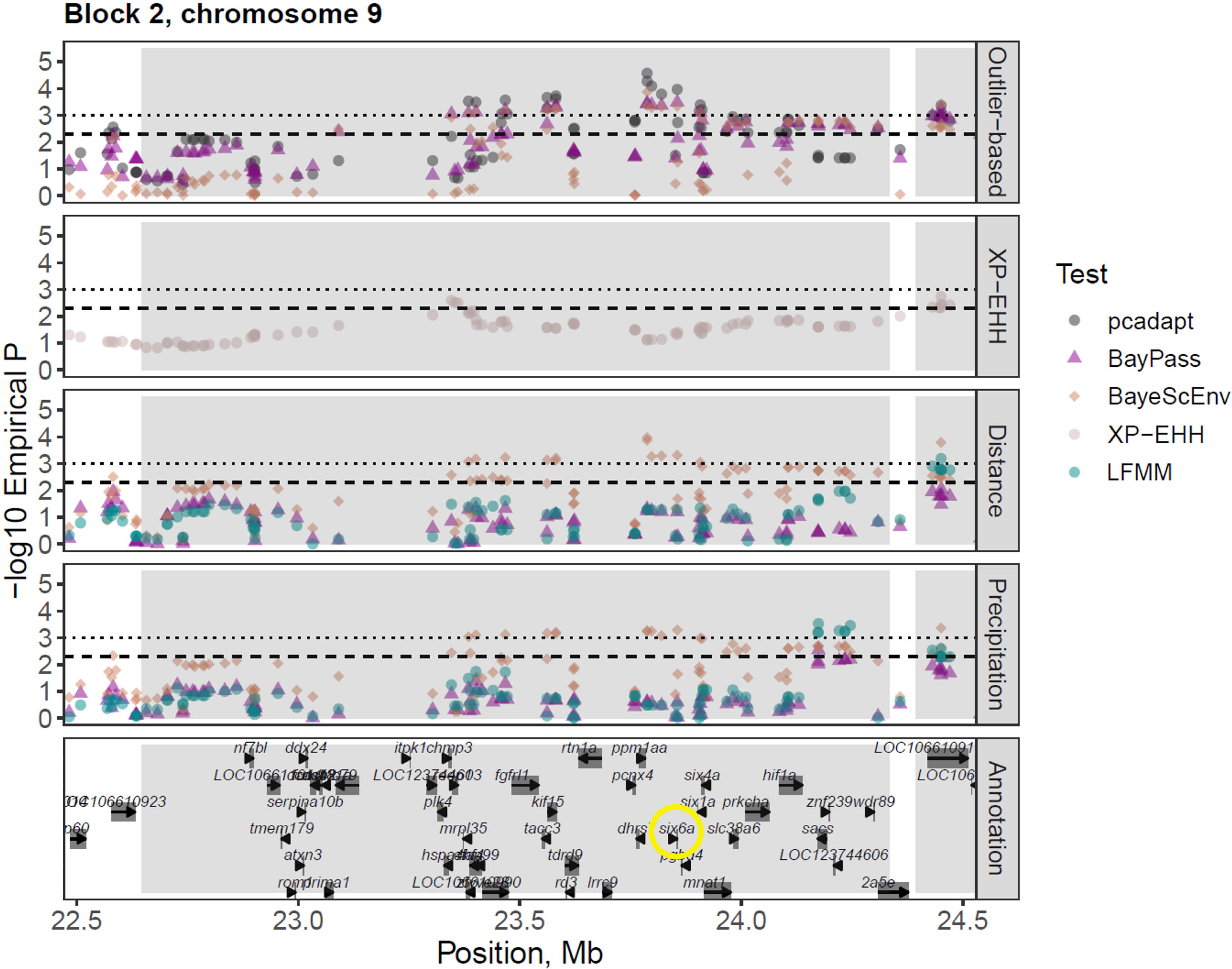
Signatures of local selection on the chromosome 9 region containing the*six6* gene (circled in yellow). The dashed and dotted lines indicate empirical p < 0.005 and p < 0.001, respectively (empirical p = SNP rank/total number of tests). The grey rectangles show the boundaries of candidate haploblocks identified in the study (Table 3). The panels from top to bottom show empirical p-values for outlier-based tests, XP-EHH test, univariate GEA analyses of distance from river mouth (also a proxy for e.g. temperature-related variation) and of the environmental variable (annual precipitation, i.e. BIO12) associated with candidate SNPs in the haploblock in both univariate and multivariate GEA analyses.

#### 3.3.5 Overlap with previously identified structural variants

We found six of the 311 unique candidate SNPs from outlier-based, univariate GEA and XP-EHH analyses to overlap with structural variants identified in Bertolotti et al. (2020): five of them were labelled as deletions, and one of them as an inversion. Two of the candidate SNPs within our candidate haploblocks were among these genomic rearrangements (both were labelled as deletions, Table 3). Our candidate haploblocks also contained more structural variants (Table S6), but they did not directly overlap with our candidate SNPs.

## 4. Discussion

Using a genome-wide SNP array, this study characterised the fine-scale population genetic structure of a large Atlantic salmon stock and identified candidate loci exhibiting signals of local selection. The putatively adaptive loci identified here partly overlapped with candidate loci previously detected in other salmonids and across the range of Atlantic salmon, providing further evidence for a recurring genetic basis of local selection across evolutionary lineages.

### 4.1 Genetic diversity and population structure

We identified four relatively distinct genetic clusters in the Tornio-Kalix River complex, consisting of the lower parts of both rivers, upper parts of both rivers, the Ängesån tributary in the Kalix River, and the upper part of the Lainio tributary in the Tornio/Torne River. This is largely in line with previous studies using small, neutral sets of markers (Jansson, 1993; Miettinen et al., 2021; Ståhl, 1981; Ståhl, 1983). When using partial redundancy analyses (pRDA) to decompose the possible drivers of population differentiation in the river system, we found a small but significant portion (6.9%) of the genetic variation to be explained by a combination of geography/environment, climate, and genetic structure (Table S4). Each set of these variables independently explained a significant proportion (a fourth) of the total explained genetic variation. The remaining fourth of the explained variation could not be uniquely attributed to any specific set of predictors (Capblancq & Forester, 2021; Capblancq, Luu, Blum, & Bazin, 2018). Although the fraction of remaining unexplained variation was very large (93.1%), these results suggest that geographic/environmental and climatic variation play a role in differentiating loci and sites in the system. The high proportion of unexplained variation could be due to genetic drift within the subpopulations (Meirmans, 2015), and/or the variables used here not capturing some important drivers of selection in the river system.

Stocking is known to cause homogenisation of salmonid stocks (Östergren et al., 2021), and could be a factor behind the lack of a stronger population structure in the Tornio-Kalix system. Large-scale stocking took place in many regions of the Tornio system from 1977 to 2002 (with offspring of adult salmon caught in the Tornio River), during and after a bottleneck that led to the near-extinction of the stock in the late 1980s (Anttila, Romakkaniemi, Kuusela, & Koski, 2008; Pruuki, 1993; Romakkaniemi et al., 2003; Romakkaniemi, 2008). This could explain why the “downstream” genetic cluster identified here extended surprisingly high up in the Tornio River (up until site To10; Figure 2b). The Ängesån and upper parts of the Lainio tributary have not been stocked as heavily as most other sites in the Tornio River, which could partly explain their genetic divergence from the other sites. However, a study using 82 SNPs compared samples from the late 1920s and 2012 and found no evidence for significant genetic changes in the Tornio or Kalix stocks over time (Östergren et al., 2021).

The Jokkfall (above site Ka2) waterfall in the Kalix River was a partial migration barrier until the construction of a fish ladder in the 1980s (Jansson, 1993), but the lack of migration obstacles in the main stem of the Tornio River (Romakkaniemi et al., 2003) could explain the relatively shallow population structure in the river system. Also, our results indicated notably high gene flow between the Tornio and Kalix Rivers. A natural explanation for this could be the bifurcation (site Ka8) that connects the two rivers, providing a natural passage for salmon from one river to enter the other. Individuals from this site appeared genetically intermediate between the upstream and downstream clusters (Figure 2), suggesting some gene flow through it (Miettinen et al., 2021). Also, the mixing of waters from the two rivers caused by the bifurcation could be thought to increase the rate of salmon originating from one river to stray to the other during spawning migration (due to limited distinction of the rivers via olfactory cues) (Miettinen et al., 2021), which is supported by tagging studies demonstrating straying to be rather common among the Tornio and Kalix (A. Romakkaniemi, unpublished data). Additionally, a small proportion of individuals sampled from downstream sites genetically resembled upstream subpopulations (Figure 2), possibly suggesting that upstream individuals may end up spawning in lower parts of the system.

Salmon from the Ängesån, upper Lainio and the other upper parts of the river system exhibited lower mean heterozygosities (similar to Jansson, 1993; Miettinen et al., 2021) and on average longer runs of homozygosity (ROH) compared to the downstream genetic cluster, potentially indicating higher levels of inbreeding in these subpopulations. This may suggest that demographic bottlenecks and thus high drift have at least partly driven the genetic divergence of these subpopulations, likely along with adaptive processes (see Bradbury et al., 2014). Future analyses of gene flow within the river system could help resolve the role of different factors in the observed population structure.

### 4.2 Genomic signatures of local adaptation

Using a combination of genotype-environment association (GEA, including redundancy analyses; RDA) analyses and outlier- and haplotype-based approaches, we found signals of putatively adaptive genetic variation related to several environmental/geographic and climatic variables, such as precipitation and distance from river mouth. Here, we only discuss genomic regions (candidate haploblocks) that contained candidates from multiple types of analyses. Many of the genes found in the candidate haploblocks are related to previously documented major life-history variation, such as the timing or length of migration (*six6*, *mc4r*, e.g. Pritchard et al., 2018), age at maturity (*six6, taar13c-like*, e.g. Barson et al., 2015; Sinclair-Waters et al., 2022) or feeding and appetite control in Atlantic salmon (*pomca, mc4r*, e.g. Kalananthan et al., 2023, 2020; Norland, Eilertsen, Rønnestad, Helvik, & Gomes, 2023). Many of the candidates also included loci previously identified as outliers among or within salmonids of different lineages (e.g. López, Cádiz, Rondeau, Koop, & Yáñez, 2021), including comparisons of wild and domesticated Atlantic salmon (e.g. López et al., 2019). These results therefore suggest a level of genomic parallelism underlying putative local adaptation across broad geographical areas and evolutionary lineages of salmonids.

In general, the allele frequencies of many candidate SNPs closest to the candidate genes reflected the upstream-downstream divergence in the river system reasonably closely. Within-river variation of the candidate genes involved in migration and maturation in particular could potentially explain previously detected life history diversity linked with population structure in this river system: smolt and adult migration timing (inferred from smolt and adult catches from the Tornio River) of salmon originating from the upper river sections has been found to differ from their downstream counterparts (Miettinen et al., 2021).

It should be noted that the north-south geography of the Tornio-Kalix River system (Figure 1), as well as the complex life history of Atlantic salmon, complicate interpretations of the drivers of putative adaptive differentiation. Population structure, geography and some environmental/climatic predictors are correlated in this river system, and therefore the drivers of selection signatures may be difficult to distinguish (Excoffier et al., 2009; Wang & Bradburd, 2014). Due to strong correlations between variables such as elevation, latitude, and annual mean temperature, we retained only distance from river mouth that can be considered a proxy for these and other relevant environmental variables (Figure S1), including migration difficulty (distance x elevation) and the “north-south gradient” that in turn reflects the harshness of environmental and climatic conditions experienced by the salmon juveniles. In addition, we performed RDA with a partial model accounting for population structure, and with a “full” model not accounting for it. These two models detected very different SNPs (Figure S5), which likely reflects that the partial model prevented the detection of many potentially adaptive environmental/climate-associated SNPs identified by the other analyses, especially on Chr 9, because of their strong association with population structure (Forester et al., 2018; Salisbury et al., 2023). Thus, we also consider the results of the full RDA model relevant (see Forester et al., 2018).

#### 4.2.1 Functional relevance of identified candidate genes

A large proportion of the signals of selection were detected in several regions on chromosome 9 (Chr 9). Our results provide additional support for a particular genomic region on this chromosome to repeatedly underlie adaptive diversification in Atlantic salmon (Pritchard et al., 2018): this candidate haploblock contained the highest number of annotated genes (n = 50, Figure 3), and included candidate SNPs identified by all analysis methods in our study. This block (hereafter “ *six6* block”) included the *six6* gene that encodes an evolutionarily conserved transcription factor and is expressed in the developing eyes, nose, brain, gill, and testis of salmon (Kurko et al., 2020; Moustakas-Verho et al., 2020). It is involved in fine- and broad-scale local adaptation and/or spatial differentiation within and among multiple lineages of Atlantic salmon (Gabián, Morán, Saura, & Carvajal-Rodríguez, 2022; Pritchard et al., 2018; Zueva et al., 2021), as well as Pacific salmonids (Andrews et al., 2023; Tigano & Russello, 2022), and is thus a strong candidate for being the selective target in the genomic region in this study.

The *six6* gene has been found to be associated with age at maturity in multiple studies of wild and aquaculture Atlantic salmon (Barson et al., 2015; Besnier et al., 2023; Jensen et al., 2022; Kess et al., 2022; Sinclair-Waters et al., 2022, 2020), and in Pacific salmonids (Waters et al., 2021; Willis et al., 2020). It has also been found to epistatically interact with another large-effect locus involved in salmon maturation (*vgll3* on Chr 25), regarding age at maturity (Besnier et al., 2023) and maximum metabolic rate (Prokkola et al., 2022). However, it is possible that instead of age at maturity, *six6* is associated with some other correlated trait in a population-specific way (e.g. size at maturity or migration timing; Barson et al., 2015, Pritchard et al., 2018, Wellband et al., 2018, Zueva et al., 2021, Kess et al., 2022). For example, allele frequencies of *six6* gene have been found to vary with migration timing of Atlantic salmon in the Atlantic and Barents-White Sea lineages (Cauwelier, Gilbey, Sampayo, Stradmeyer, & Middlemas, 2018; Pritchard et al., 2018), and along an elevation gradient in the North American lineage in the Miramichi River (Wellband et al., 2018). Similar to the Tornio River (Miettinen et al., 2021), salmon spawning in the upper river sections enter the Miramichi earlier in the migration season (Chaput, Douglas, & Hayward, 2016) and had the highest alternate *six6* allele frequencies among tributaries in the system (Wellband et al., 2018). Interestingly, Cauwelier et al. (2018) also found an upstream-downstream genetic population structuring across multiple salmon rivers in Scotland, with the *six6* genomic region being associated with migration timing differences in some of the rivers (Cauwelier et al., 2018). We also observed upstream individuals (except for the upper Lainio subpopulation) in the Tornio-Kalix system to exhibit a higher frequency of the “early migration timing” allele of *six6* as in Cauwelier et al. (2018) and Pritchard et al. (2018). This implies that *six6* may be associated with migration timing in four different evolutionary lineages of Atlantic salmon (Baltic, Barents-White Sea, Scottish/North Atlantic and North American).

Genetic variation in the *six6* block was associated with annual precipitation, precipitation of the wettest quarter (i.e. precipitation in the summer), distance from river mouth, and water velocity. This is similar to a *six6* association with winter precipitation in the Miramichi River (Wellband et al., 2018). Precipitation has been described as a potential driver of adaptive genetic differentiation among salmonid populations (e.g. Bourret, Kent, Lien, & Bernatchez, 2013; Hecht, Matala, Hess, & Narum, 2015). Within-river variation in precipitation could possibly be involved in e.g. the timing of migrations and/or in fish size through its effects on water flow (Wellband et al., 2018). Precipitation also affects run-off and therefore likely water colour, visibility and turbidity, which could be connected to the suggested of roles of *six6* in spatial cognition, navigation, foraging and spawning site selection (Pritchard et al., 2018, Moustakas-Verho et al., 2020), and to this gene being involved in adaptive, vision-related responses to variation in habitats experienced by salmonids (Tigano & Russello, 2022). Therefore, variation in *six6* could be an adaptive response to differences in visual habitats among the sites (Tigano & Russello, 2022). Finally, *six6* has been found to be associated with diet acquisition strategies of Atlantic salmon in the marine environment (Aykanat et al., 2020). We therefore argue that *six6* may have multiple adaptive roles for salmon in this river system, and that some of them may be life stage-specific (Aykanat et al., 2020), i.e. not only associated e.g. with the juvenile and spawning environment. The *six6* variation we observed may also reflect differential selective pressures experienced by the subpopulations regarding e.g. food intake, age at maturity and/or migration timing.

Three other candidate blocks also contained genes involved in reproductive timing (Table S5). For example, *taar13c-like* was associated with age at maturity in a study of wild Atlantic salmon populations from the Atlantic lineage (Sinclair-Waters et al., 2022), and its paralogue was a candidate under putative selection among salmon originating from freshwater lakes and the Atlantic Ocean (Zueva, Lumme, Veselov, Kent, & Primmer, 2018). *Grm4* was associated with migratory behaviour (resident vs. migratory life history) of brown trout in Finland (Lemopoulos, Uusi-Heikkilä, Huusko, Vasemägi, & Vainikka, 2018). It belongs to the metabotropic glutamate receptors that have been found to be under differential selection or methylation between resident and migratory rainbow trout ecotypes (Baerwald et al., 2016; Hale, Thrower, Berntson, Miller, & Nichols, 2013).

Morphology- or growth-related genes (in 11/18 blocks) included *bdkrb2, acv2rb and gpr4* that are associated with (muscle) growth in Atlantic salmon or rainbow trout (Barría et al., 2019; Phelps, Jaffe, & Bradley, 2013; Weber, Ma, Birkett, & Cleveland, 2022; Yoshida, Lhorente, Carvalheiro, & Yáñez, 2017; Yoshida & Yáñez, 2022). Diet- or feeding-associated genes (in 7/18 blocks) included *pomca* (*proopiomelanocortin a precursor*, hypothalamic neuropeptide) and *mc4r (melanocortin receptor 4)*, both in the melanocortin system and involved in appetite control and feeding of Atlantic salmon (e.g. Kalananthan et al., 2023, 2020; Norland et al., 2023). In addition to appetite control and feeding, *mc4r* has a key role in energy homeostasis and somatic growth/weight gain in fish and other vertebrates (Krashes, Lowell, & Garfield, 2016; Metz, Peters, & Flik, 2006; Khalil et al., 2023). This gene or a homologous region has been found to be associated with migration distance or difficulty in salmonids (Moore et al., 2017; Pritchard et al., 2018). As individuals from the upstream reaches of the Tornio-Kalix migrate earlier in the spawning migration season (Miettinen et al., 2021), they may be fasting for a longer time than their downstream counterparts, because Atlantic salmon are not thought to feed during their spawning migration (Jonsson, Jonsson, & Hansen, 1997). To add, a 3110bp deletion in the *pomca* haploblock was tagging the gene *capn1-like* (*calpain1 catalytic subunit-like*) whose expression is triggered following starvation in rainbow trout juveniles (Salem, Nath, Rexroad, Killefer, & Yao, 2005). Thus, e.g. appetite, growth and starvation resistance could be expected to adaptively vary in salmon populations in response to migration duration and difficulty, which could possibly explain our results for the genes mentioned here.

Interestingly, almost all candidate blocks (16 out of 18) contained genes associated with temperature/thermal adaptation, such as *hspa4l* that is heavily involved in heat stress responses in rainbow trout (Huang, Li, Liu, Kang, & Wang, 2018; X. Liu et al., 2021), and the *exoc5* and *farp1* genes that are associated with cold adaptation in fish (Bilyk, Zhuang, & Papetti, 2023; Sun, Huang, Kong, Wang, & Kang, 2022). Many of the temperature-related genes were on blocks that were associated with e.g. distance from river mouth and/or precipitation (i.e. variables strongly correlated with temperature and elevation; Figure S2), suggesting these associations could reflect the potential role of these genes in thermal adaptation in this system.

We found candidate genes associated with adaptation to hypoxia in many blocks (in 7/18 blocks). Hypoxia in the river system can be assumed to increase with elevation from sea level, resulting from a lower barometric pressure causing a decrease in oxygen levels. Distance from river mouth was very strongly correlated with elevation, and therefore candidate blocks associated with it may include particularly relevant candidate genes involved in altitude and hypoxia adaptation, such as the major regulators of hypoxia responses *hif1a* and *eif5* (Braz-Mota & Almeida-Val, 2021; Tariq, Ito, Ishfaq, Bradshaw, & Yoshida, 2016), and *rock2* (Chen et al., 2020).

Water chemistry has been recognised as a potential selective agent driving local adaptation in Atlantic salmon (Bourret et al., 2013). As precipitation affects run-off and therefore likely water chemistry too, alkalinity- or acidity-related candidate genes (in total in 5/18 blocks), such as *bdkbr1* and *bdkrb2* that were in a block associated with precipitation, could be thought to be involved with local adaptation to the water chemistry of the juvenile environment. On the other hand, salinity-associated genes were found on multiple blocks (in 10/18), and genes such as *odc1* (e.g. Whitehead, Roach, Zhang, & Galvez, 2011) and *arf1* (Gao, Xu, & Xu, 2021) could possibly indicate differential selective pressures experienced by salmon from different parts of the river system during their marine phase in the Baltic Sea that has a strong salinity gradient. Future studies examining possible subpopulation-specific migration routes at sea and/or water chemistry of the juvenile/spawning environment could improve our understanding of the selective pressures faced by salmon originating from different sites and at different life stages.

In addition to *six6* (see above), we found many genes associated with light and/or vision (in 6/18 blocks), situated on blocks associated with precipitation, water velocity, water depth, bottom substrate and/or underwater vegetation. Variation in these variables likely affects e.g. water colour, turbidity, visibility and general light conditions. Genes such as *exoc5* and *otx2b* are associated with eye development (Lobo et al., 2017; Kurko et al., 2020), while *rgra* codes a vision-related intracellular signalling protein and may be involved with adaptation to differences in light conditions (e.g. Liu et al., 2020; K. Wang et al., 2019; Perez et al. 2021). *Rp1l1* has been suggested to facilitate visual adaptation of benthic ectomorphs of cichlids to darkness (Malinsky et al., 2015), and was found here on a candidate block with overall eight genes possibly associated with vision or light. Differences in vision-related genes have also been observed among pink salmon populations, with the potential adaptive explanation being the transitioning from planktivorous to piscivorous diets during the salmon life cycle, which requires developing e.g. night vision to chase prey (Christensen et al., 2021). Therefore, the vision- or light-related genes discussed here could be adaptively important in different parts of the life cycle of salmon.

#### 4.2.2 Limitations

We note that loci under weaker selection might not have been detected by our approach of accounting only for candidate genomic regions overlapping across multiple types of analyses. Furthermore, some of the candidate regions identified here could also be related to gene regulation, and/or be a result of historical selection and might thus not contribute to phenotypic variation in the current environment (reviewed by Waples et al., 2022). Therefore, complementary analyses and experimental manipulations are needed to better understand how the candidate loci may be involved in local adaptation.

### 4.3 Management implications

We identified putatively adaptive differentiation at the functional genetic level in the Tornio-Kalix system, indicating that adaptive processes have a role in shaping the population structure previously studied only with few neutral markers. The adaptive genetic diversity demonstrated to exist within the river system should be monitored and considered in fisheries management (as recommended in general by e.g. Miettinen et al., 2021; Bernatchez et al., 2017). The importance of population substructuring in Baltic salmon was also recently highlighted by a modelling study demonstrating that extinction of local subpopulations (at the river level) can reduce the genetic diversity of the surviving subpopulations, suggesting that the loss of genetic diversity within Baltic salmon has probably been overlooked, and that the Tornio-Kalix stock (and its substructure) is crucial for the genetic diversity of Baltic salmon as a whole (Kurland, Ryman, Hössjer, & Laikre, 2023). This further underscores the importance of preserving local subpopulations, and provides support that salmon should be managed at the subpopulation level (see Pritchard et al., 2018; Vähä, Erkinaro, Falkegård, Orell, & Niemelä, 2017; Vähä, Erkinaro, Niemelä, & Primmer, 2007, 2008).

Preserving adaptive genetic diversity can protect a wide range of life history variation that can be essential for the evolutionary and harvesting potential of populations (e.g. Carvalho et al., 2023). Although most of the genetic variation relevant to management and conservation is likely polygenic, identifying adaptive candidate loci is the first step to developing e.g. genetic assays that can be used in monitoring and guiding management strategies to help preserve variation at these loci (Hohenlohe et al., 2021). Information from this study can thus be used to develop genetic markers for monitoring changes in allele frequencies of the ecologically relevant candidate adaptive loci identified here, and for genetic stock identification of Baltic salmon catches to identify them to their subpopulations of origin. This can in turn help examine the influence of human-induced selective pressures (such as climate change and fishing practices) on the subpopulations and candidate adaptive loci across time and space (both in the future and in the past; e.g. Kardos & Shafer, 2018).

Allele frequencies of large-effect loci in Atlantic salmon can respond to selection rapidly, demonstrated e.g. by the substantial drop in the frequency of the *six6* allele associated with older age at maturity after a dam construction in River Eira in Norway (Jensen et al., 2022). This highlights that such loci can be suitable diagnostic markers for understanding rapid evolutionary changes in wild populations (see also Czorlich, Aykanat, Erkinaro, Orell, & Primmer, 2022; Czorlich, Aykanat, Erkinaro, Orell, & Primmer, 2018). Furthermore, an improved understanding of the environmental drivers of local adaptation may allow predicting future trajectories of salmon stocks and identifying particularly vulnerable populations under different scenarios of environmental change (e.g. Capblancq, Fitzpatrick, Bay, Exposito-Alonso, & Keller, 2020; Capblancq & Forester, 2021; Layton et al., 2021).

Finally, the relatively weak genetic structure of the Tornio-Kalix salmon stock may also indicate a possible homogenising effect of past stocking activities on its genetic diversity. This further underlines the need for caution when considering future stocking in this or other river systems, to account for the potential within-river locally adaptive differences (see Östergren et al., 2021).

### 4.4 Conclusions

Our study elucidated the genetic population substructuring and identified multiple strong candidates for loci under fine-scale local selection in the largest Baltic salmon stock. These candidates are involved in life history variation of ecologically important traits in multiple salmonid lineages. Most notably, our results provide further support for the *six6* genomic region to be adaptively important over the global range of salmonids.

## Author contributions

Conceptualisation: VLP, AM, CRP

Data curation: AM, JD

Formal analysis: AM, EPA

Funding acquisition: AM, VLP, AR, SP, JÖ, JD, CRP

Methodology: VLP, AM

Project administration: AM

Resources: AR, CRP

Supervision: VLP, CRP

Visualisation: AM

Writing (original draft): AM

Writing (review & editing): All authors

## Acknowledgements

We are grateful to field workers and other staff for sample provision and assistance with research material (Rauno Hokki, Kari Pulkkinen, Markku Kilpala), including laboratory assistance (Linda Söderberg, Iikki Donner, Morgan Frapine). We acknowledge CIGENE (Centre for Integrative Genetics, Norwegian University of Life Sciences, Norway) for genotyping our samples, and CSC (IT Center for Science, Finland) for computational resources. We thank the EvolConGen research group and UHI Institute for Biodiversity & Freshwater Conservation for discussions.

This study was funded by Tornio/Torne River fishing license revenues, The Swedish Agency for Marine and Water Management, The Swedish Research Council Formas (Grant/Award 2013-1288 to JÖ), The Finnish Society of Sciences and Letters (grant to VLP), Societas pro Fauna et Flora Fennica, The Betty Väänänen Fund from The Kuopio Naturalists’ Society (KLYY, Betty Väänäsen rahasto), The Raija and Ossi Tuuliainen Foundation (Raija ja Ossi Tuuliaisen Säätiö), The Baltic Sea Fund from The Finnish Foundation for Nature Conservation (Suomen Luonnonsuojelun Säätiö), the Alfred Kordelin Foundation (Alfred Kordelinin säätiö), and the LUOVA doctoral programme (University of Helsinki) (grants and funding to AM), the Finnish Society of Sciences and Letters (to VLP), the University of Helsinki, and the European Union (ERC, FishLEGs, 101054307) (to CRP).

