## Supplementary materials (Tables S1-S6 and Figures S1-S6) for "Loci associated with maturation, migration and appetite control are linked with signals of fine-scale local selection in a large Atlantic salmon population"

**Table S1.** Sample site information with summary statistics for genetic diversity. Hexp = expected heterozygosity, Hobs = observed heterozygosity, ROH (KBAVG) = mean length of ROH per site.

| Site | # locations | Genetic cluster | Genotyped (n) | Successful^b^ (n) | After filtering (n) | H_exp_ | H_obs_ | F_IS_ | ROH (KBAVG) | Previous name^c^ |
| --- | --- | --- | --- | --- | --- | --- | --- | --- | --- | --- |
| Ka1 | 4 | Lower TK^a^ | 20 | 20 | 20 | 0.356 | 0.358 | 0.016 | 6838.6 | K1 |
| Ka2 | 2 | Lower TK | 21 | 21 | 21 | 0.357 | 0.365 | -0.001 | 5630.1 | K2 |
| Ka3 | 3 | Upper TK | 20 | 20 | 20 | 0.342 | 0.353 | -0.005 | 6060.2 | K3 |
| Ka4 | 2 | Upper TK | 20 | 20 | 19 | 0.332 | 0.337 | 0.011 | 9246.2 | K4 |
| Ka5 | 3 | Upper TK | 20 | 20 | 20 | 0.338 | 0.348 | -0.003 | 6994.4 | K5 |
| Ka6 | 1 | Ängesån | 17 | 17 | 16 | 0.342 | 0.356 | -0.012 | 7466.2 | K6 |
| Ka7 | 1 | Ängesån | 17 | 17 | 17 | 0.335 | 0.343 | 0.007 | 10917.9 | K7 |
| Ka8 | 2 | Lower TK | 20 | 20 | 20 | 0.352 | 0.362 | -0.005 | 5053.9 | K8 |
| To1 | 4 | Lower TK | 20 | 20 | 19 | 0.354 | 0.365 | -0.003 | 6336.3 | T1 |
| To2 | 4 | Lower TK | 20 | 17 | 17 | 0.355 | 0.380 | -0.031 | 3219.5 | T2 |
| To3 | 3 | Lower TK | 20 | 20 | 20 | 0.355 | 0.366 | -0.005 | 4322.6 | T3 |
| To4 | 4 | Upper TK | 20 | 20 | 20 | 0.343 | 0.357 | -0.013 | 5360.4 | T4 |
| To5 | 2 | Lower TK | 20 | 20 | 20 | 0.352 | 0.363 | -0.003 | 6305.1 | T5 |
| To7 | 2 | Lainio | 20 | 20 | 20 | 0.334 | 0.345 | -0.007 | 9997.7 | T6 |
| To8 | 4 | Lower TK | 20 | 19 | 19 | 0.352 | 0.364 | -0.007 | 4483.6 | T7 |
| To9 | 2 | Lower TK | 20 | 18 | 18 | 0.352 | 0.369 | -0.016 | 5934.6 |  |
| To10 | 2 | Lower TK | 20 | 19 | 19 | 0.351 | 0.365 | -0.010 | 5395 |  |
| To11 | 3 | Upper TK | 21 | 21 | 17 | 0.341 | 0.358 | -0.019 | 5102.1 |  |
| To12 | 4 | Upper TK | 20 | 20 | 19 | 0.344 | 0.357 | -0.009 | 6525.4 | T8 |
| Total | 52 |  | 376 | 369 | 361 |  |  |  |  |  |

^a^ TK = Tornio-Kalix

^b^ Passed genotyping quality control

^c^ If site was included in Miettinen et al. (2021)

**Table S2.** Information of the 26 environmental, geographic and bioclimatic variables available for the sampling locations in our dataset.

| Variable | Measured in | Note | Variable class | Category of measurement unit | Obtained from | Retained in analyses |
| --- | --- | --- | --- | --- | --- | --- |
| Distance from river mouth | km |  | Geographic | Continuous | Luke/SLU | Yes |
| Elevation | m | Highly correlated with distance from river mouth | Geographic | Continuous | Google Earth | No |
| Latitude |  | Highly correlated with distance from river mouth | Geographic | Continuous | Google Earth | No |
| Mean depth | cm |  | Environmental | Continuous | Luke/SLU | Yes |
| Mean velocity of water flow | m/s | Classified as “calm” (< 0.2), “medium” (0.2-0.7) or “rapid” (> 0.7) | Environmental | Categorical | Luke/SLU | Yes |
| Dominant bottom substrate | particle diameter size in cm | Classified as “fine” (< 2), “stones” (2-10), “rocks” (10-30), “large rocks” (> 30) | Environmental | Categorical | Luke/SLU | Yes |
| Coverage of underwater vegetation | % | Classified as “little”, “medium”, “abundant” | Environmental | Categorical | Luke/SLU | Yes |
| Annual mean temperature (BIO1) |  | Highly correlated with distance from river mouth | Bioclimatic | Continuous | WorldClim^a^ | No |
| Mean diurnal range (BIO2) |  | Mean of monthly (max temp-min temp) | Bioclimatic | Continuous | WorldClim^a^ | No |
| Isothermality (BIO3) |  | (BIO2/BIO7)×100 | Bioclimatic | Continuous | WorldClim^a^ | Yes |
| Temperature seasonality (BIO4) |  | Standard deviation ×100 | Bioclimatic | Continuous | WorldClim^a^ | No |
| Maximum temperature of warmest month (BIO5) |  |  | Bioclimatic | Continuous | WorldClim^a^ | No |
| Minimum temperature of coldest month (BIO6) |  |  | Bioclimatic | Continuous | WorldClim^a^ | No |
| Temperature annual range (BIO7) |  | BIO5-BIO6 | Bioclimatic | Continuous | WorldClim^a^ | Yes |
| Mean temperature of wettest quarter (BIO8) |  |  | Bioclimatic | Continuous | WorldClim^a^ | Yes |
| Mean temperature of driest quarter (BIO9) |  |  | Bioclimatic | Continuous | WorldClim^a^ | No |
| Mean temperature of warmest quarter (BIO10) |  |  | Bioclimatic | Continuous | WorldClim^a^ | No |
| Mean temperature of coldest quarter (BIO11) |  |  | Bioclimatic | Continuous | WorldClim^a^ | No |
| Annual precipitation (BIO12) |  |  | Bioclimatic | Continuous | WorldClim^a^ | Yes |
| Precipitation of wettest month (BIO13) |  |  | Bioclimatic | Continuous | WorldClim^a^ | No |
| Precipitation of driest month (BIO14) |  |  | Bioclimatic | Continuous | WorldClim^a^ | No |
| Precipitation seasonality (BIO15) |  | Coefficient of variation | Bioclimatic | Continuous | WorldClim^a^ | No |
| Precipitation of the wettest quarter (BIO16) |  |  | Bioclimatic | Continuous | WorldClim^a^ | Yes |
| Precipitation of the driest quarter (BIO17) |  |  | Bioclimatic | Continuous | WorldClim^a^ | No |
| Precipitation of the warmest quarter (BIO18) |  |  | Bioclimatic | Continuous | WorldClim^a^ | No |
| Precipitation of the coldest quarter (BIO19) |  |  | Bioclimatic | Continuous | WorldClim^a^ | No |

^a^ Obtained using a resolution of 2.5 minutes of a degree.

**Table S3.** Pairwise F_ST_ estimates among the 19 sampling sites.

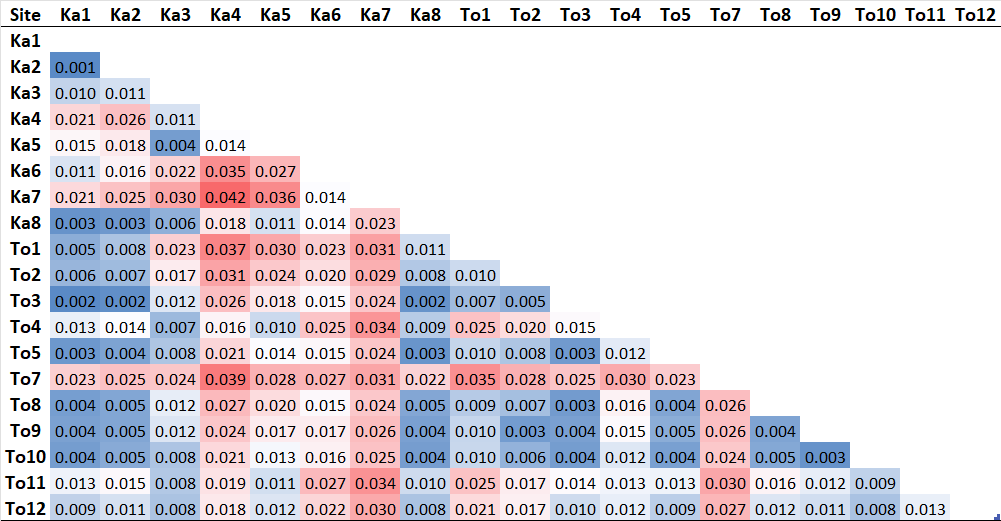

**Table S4.** Variance partitioning by partial RDA models using different sets of variables (“climatic” = “Pure climate”, “environmental/geographic” = “Pure environment/geography”, and “genetic structure” = “Pure structure”) to determine the proportions of genetic variation explained by them, and by the “full RDA” model (i.e. a model that included all of the sets of variables, without conditioning any of them). The proportion of explainable variance (Prop. of expl. var.) is the total constrained variation explained by the full model.

| (Partial) RDA models | Variance | Adj. R^2^ | F | Pr(>F) | Prop. of expl. var. | Prop. of total variance |
| --- | --- | --- | --- | --- | --- | --- |
| Full model: F ~ clim. + env. + struct. | 881.9 | 0.0297 | 1.846 | 0.001 | 1 | 0.069 |
| Pure climate: F ~ clim. \| (env. + struct.) | 218.1 | 0.0026 | 1.187 | 0.001 | 0.247 | 0.017 |
| Pure environment/geography: F ~ env. \| (clim. + struct.) | 224.3 | 0.003 | 1.221 | 0.001 | 0.254 | 0.018 |
| Pure structure: F ~ struct. \| (clim. + env.) | 221.9 | 0.0084 | 2.013 | 0.001 | 0.252 | 0.017 |
| Confounded | 217.6 |  |  |  | 0.247 | 0.017 |
| Total unexplained | 11869.1 |  |  |  |  | 0.931 |
| Total variance | 12751 |  |  |  |  | 1 |

**Table S5.** Candidate haploblocks, their associations with environmental variables used in this study, and candidate genes in the blocks found to be potentially involved with local adaptation to environmental variation in other studies of fish.

| **Haploblock/ Chromosome** | **Environmental association** | **Candidate genes implicated in adaptation to environmental variation, and their functional relevance** | | | | | | | | | | |
| --- | --- | --- | --- | --- | --- | --- | --- | --- | --- | --- | --- | --- |
|  |  | **Temperature** | **Salinity** | **Acidity/ alkalinity** | **Morphology and growth** | **Hypoxia** | **Infection** | **Light/**  **vision** | **Feeding/diet** | **Reproductive timing** | **Other** | **References** |
| Block 1 / 9 | Distance from river mouth | *exoc5* |  |  |  |  |  | *exoc5, otx2b* |  |  | *ap5m1, otx2b* | [1–4] |
| Block 2 / 9 | Precipitation (BIO12, BIO16), distance from river mouth, water velocity | *atxn3, serpina10b, plk4, hspa4l, rnf4, tacc3, kif15, mnat1* | *itpk1, dhrs7, six4a, slc38a6* | *tacc3, tdrd9, rtn1a* | *fgfrl1, six1a* | *mrpl35, kif15, prkcha, hif1a* | *asb2, itpk1, reep1, mrpl35, ppm1aa* | *rom1* | *six1a* | *lrrc9, pcnx4, six6a* | *kif15, tdrd9, rtn1a, dhrs7, ppm1aa, pgbd4, six6a* | [5–34] |
| Block 3 / 9 | Precipitation (BIO16), isothermality, distance from river mouth, water velocity, water depth | *eif5, plcb1, tmx1, cdc5l, clic5a, xkr6* | *kcnh5, ckb (kcrt), mark3, tnfaip2, cdc5l, clic5a, xkr6* |  | *lbh* | *eif5, plcb1* | *bag5* | *ahnak* | *lbh, hao1* |  | *eif5, hao1, cdc5l, supt3h* | [21,32,35–51] |
| Block 4 / 9 | Distance from river mouth, water depth | *eif4b, ampm1 (metap1)* | *atp10d, odc1, rock2, apoba* | *eif4b, gabra2, atp10d, lamtor3, kcnf1* | *sobp, pdss2, gdf6* | *enpep, rock2, apoba* |  |  | *lingo2, gabra1* |  | *gabra1, gabra2, gabrb1, bend3, gdf6, rrh, commd8, lamtor3, odc1, apoba* | [3,12,21,34,52–69] |
| Block 5 / 9 | Precipitation (BIO16) | *spred1* |  |  | *spred1* |  |  |  |  |  |  | [70] |
| Block 6 / 9 | Precipitation (BIO16), isothermality | *ptpn21, gpr4, bdkrb2* | *ptpn21, bdkrb1, bdkrb2* | *bdkrb1, bdkrb2* | *gpr4, bdkrb2* |  |  |  |  |  | *spata7, bdkrb2* | [53,71–75] |
| Block 7 / 9 | Precipitation (BIO16) |  | *Il17a, il17f, ccdc88c* |  | *stmn4* |  | *Il17a, il17f* |  |  |  |  | [58,76–78] |
| Block 8 / 9 | Precipitation (BIO16), isothermality | *capn1, hyou1, ephx2, dnajc27, adrb3, pygl, ninein* | *capn1, slc22a6, nhsl1, angptl7, adcy8, trim9, map3k5* |  | *tpo* | *map3k5* |  | *rp1l1, znf513, plaat1, ephx2, slc22a6, adcy8, prox1, trim9* | *pomca, capn1* |  | *pomca, prox1, capn1* | [32,56,79–103] |
| Block 9 / 13 | Precipitation (BIO16) | *ppp1r12b, grm4* |  |  |  |  |  |  |  | *grm4 (mglur4)* | *grm4 (mglur4), syt2* | [104–108] |
| Block 10 / 13 | Isothermality |  |  |  |  |  |  |  |  |  | *dyrk1aa* | [148] |
| Block 11 / 14 | Precipitation (BIO16) | *mc4r* | *gad1* |  | *rnf152* |  | *c5orf22, cdh6* |  | *mc4r* |  | *rnf31, mc4r* | [29,57,109–114] |
| Block 12 / 14 | Distance from river mouth, temperature annual range | *wnt9a* | *arf1* |  | *wnt3a* |  |  |  |  |  |  | [115–117] |
| Block 13 / 14 | Isothermality | *abca4, acvr2b, trim35* |  |  | *col11a1, acvr2b* |  |  | *abca4* | *arhgap29* | *acvr2b* |  | [70,118–122] |
| Block 14 / 17 | Distance from river mouth | *aldh1l2, etnk1* |  |  |  | *samm50, sox5* |  |  |  |  | *samm50* | [10,149–152] |
| Block 15 / 18 | Bottom substrate, underwater vegetation |  | *eif3l* | *cdh23* | *smoc2, wdr11* | *plpp4, fgfr2* | *vsir (vista)* | *rgr* |  |  |  | [123–131] |
| Block 16 / 20 | Isothermality | *selenom* | *inpp5j* |  |  |  |  |  | *stmnl1, mat1a* |  | *smtnl1* | [121,133–134, 136–137] |
| Block 17 / 21 | Precipitation (BIO16) | *farp1* |  |  |  |  |  |  |  | *taar13c-like* | *taar13c-like* | [14,138,139] |
| Block 18 / 28 | Water depth | *h3f3a* |  |  |  | *ppp1r3c* |  |  |  |  |  | [134,140] |

**Table S6.** Candidate haploblocks (i.e. blocks sharing candidate loci from at least two different types of genome scans for selection) identified in this study. This table shows information about the chromosome, position and length of each haploblock, the types of scans that identified candidates in each block, the environmental variables associated with the blocks (GEA; associations identified by both univariate and multivariate GEA are underlined), and the number of structural variants (SVs) identified by Bertolotti et al. (2020).

| Block # | Chr | Start (bp) | End (bp) | Length (Mb) | Type | GEA^a^ |  | SVs |
| --- | --- | --- | --- | --- | --- | --- | --- | --- |
| 1 | 9 | 22041594 | 22229310 | 0.19 | RDA, XP-EHH | Distance |  | 1 deletion |
| 2 | 9 | 22646469 | 24333545 | 1.69 | Outlier, GEA (both), XP-EHH | BIO12, BIO16, distance, velocity |  | 6 deletions |
| 3 | 9 | 24394915 | 25479569 | 1.08 | Outlier, RDA (both), XP-EHH | BIO3, BIO16, depth, distance, velocity |  | 13 deletions |
| 4 | 9 | 25544576 | 26572371 | 1.03 | Outlier, GEA (both), XP-EHH | Depth, distance |  | 8 deletions, 1 duplication |
| 5 | 9 | 28118150 | 28415589 | 0.30 | Outlier, RDA (both), XP-EHH | BIO16 |  | 1 deletion |
| 6 | 9 | 28487961 | 28693554 | 0.21 | Outlier, GEA (both), XP-EHH | BIO3, BIO16 |  | 1 deletion |
| 7 | 9 | 28708901 | 28998334 | 0.29 | Outlier, GEA (both) | BIO16 |  | 2 deletions |
| 8 | 9 | 44000579 | 46193652 | 2.19 | Outlier, RDA | BIO3, BIO16 |  | 13 deletions, 1 duplication |
| 9 | 13 | 33821031 | 34113068 | 0.29 | Outlier, RDA | BIO16 |  | 4 deletions |
| 10 | 13 | 72834066 | 73244495 | 0.41 | RDA, XP-EHH | BIO3 |  | 3 deletions |
| 11 | 14 | 24272662 | 24610951 | 0.34 | Outlier, RDA | BIO16 |  | 1 deletion |
| 12 | 14 | 31661331 | 31822502 | 0.16 | RDA (both), XP-EHH | BIO7, distance |  | None |
| 13 | 14 | 47642849 | 48234437 | 0.59 | RDA, XP-EHH | BIO3 |  | 2 deletions |
| 14 | 17 | 56717526 | 56866325 | 0.15 | RDA, XP-EHH | Distance |  | None |
| 15 | 18 | 1235854 | 3603311 | 2.37 | GEA (both), XP-EHH | Bottom substrate, vegetation |  | 3 deletions, 2 duplications |
| 16 | 20 | 27662780 | 27760237 | 0.10 | Outlier, RDA | BIO3 |  | None |
| 17 | 21 | 49415626 | 49962816 | 0.55 | Outlier, GEA (both) | BIO16 |  | 6 deletions |
| 18 | 28 | 6645841 | 6691045 | 0.05 | Outlier, RDA, XP-EHH | Depth |  | 1 deletion |

^a^ BIO3 = Isothermality, BIO7 = Temperature annual range, BIO12 = Annual precipitation, BIO16 = Precipitation of wettest quarter.

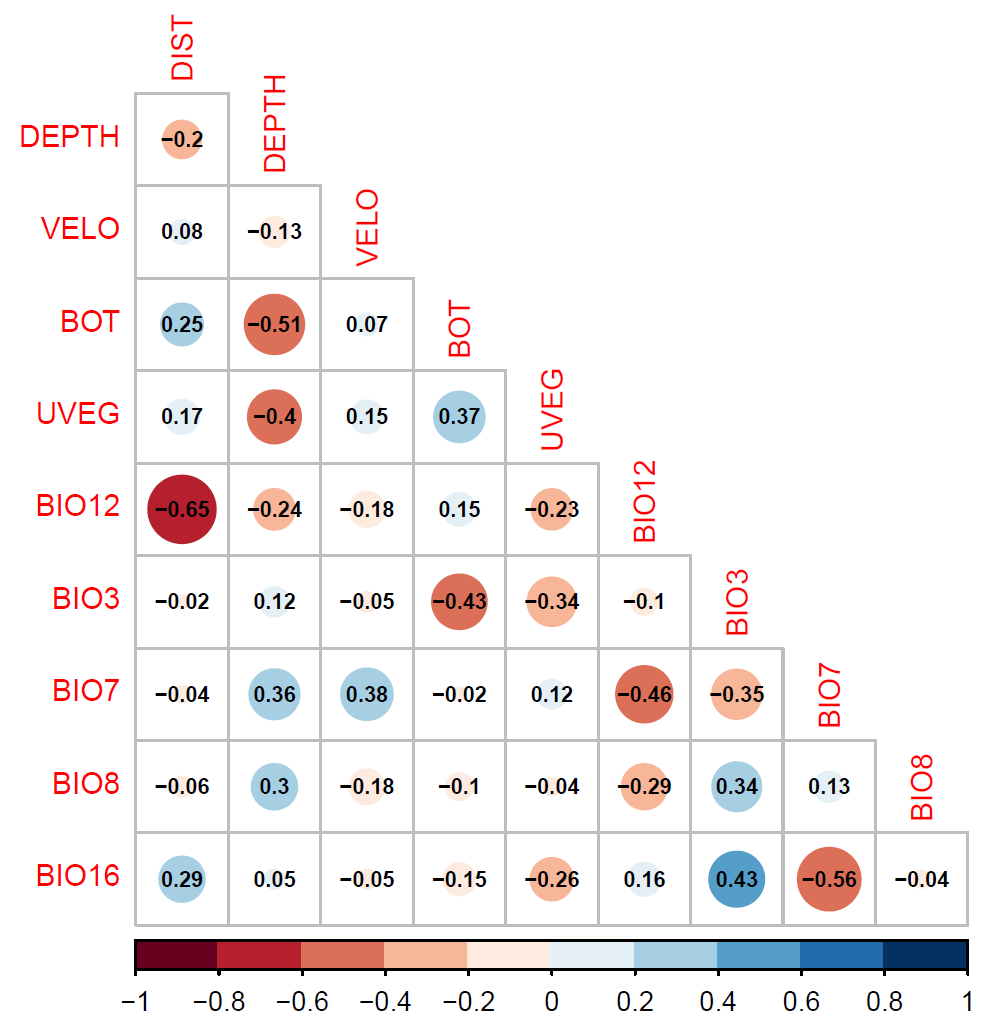

**Figure S1.** Correlation plot of the ten retained geographic, environmental and climatic variables. Values indicate Pearson correlation coefficients, and the colours and sizes of the circles the direction and strength of correlation among variables. DIST = distance from river mouth, DEPTH = mean depth, VELO = mean velocity, BOT = bottom substrate, UVEG = underwater vegetation, BIO12 = annual precipitation, BIO3 = isothermality, BIO7 = temperature annual range, BIO8 = mean temperature of the wettest quarter, BIO16 = precipitation of the wettest quarter.

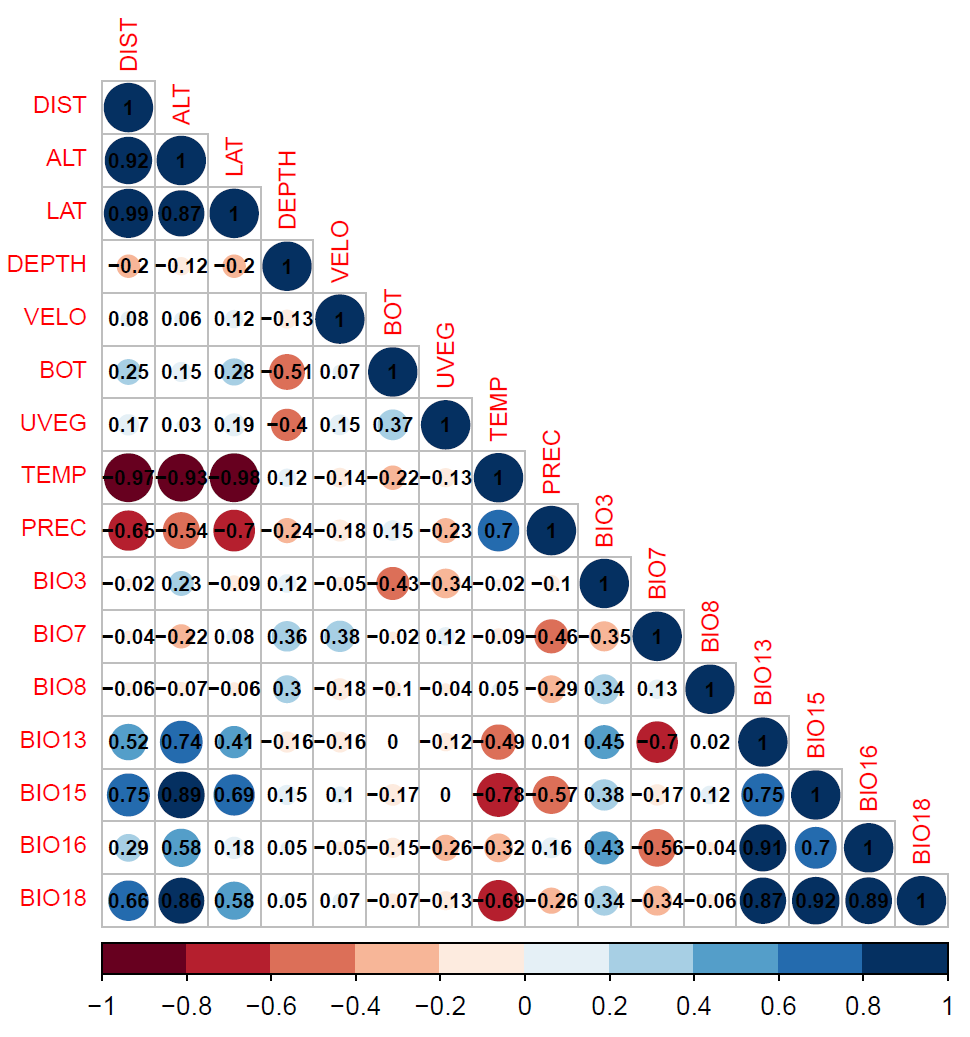

**Figure S2.** Correlation plot of all the geographic and environmental variables, and a subset of the bioclimatic variables, for which information was available for the sampling locations in our dataset.

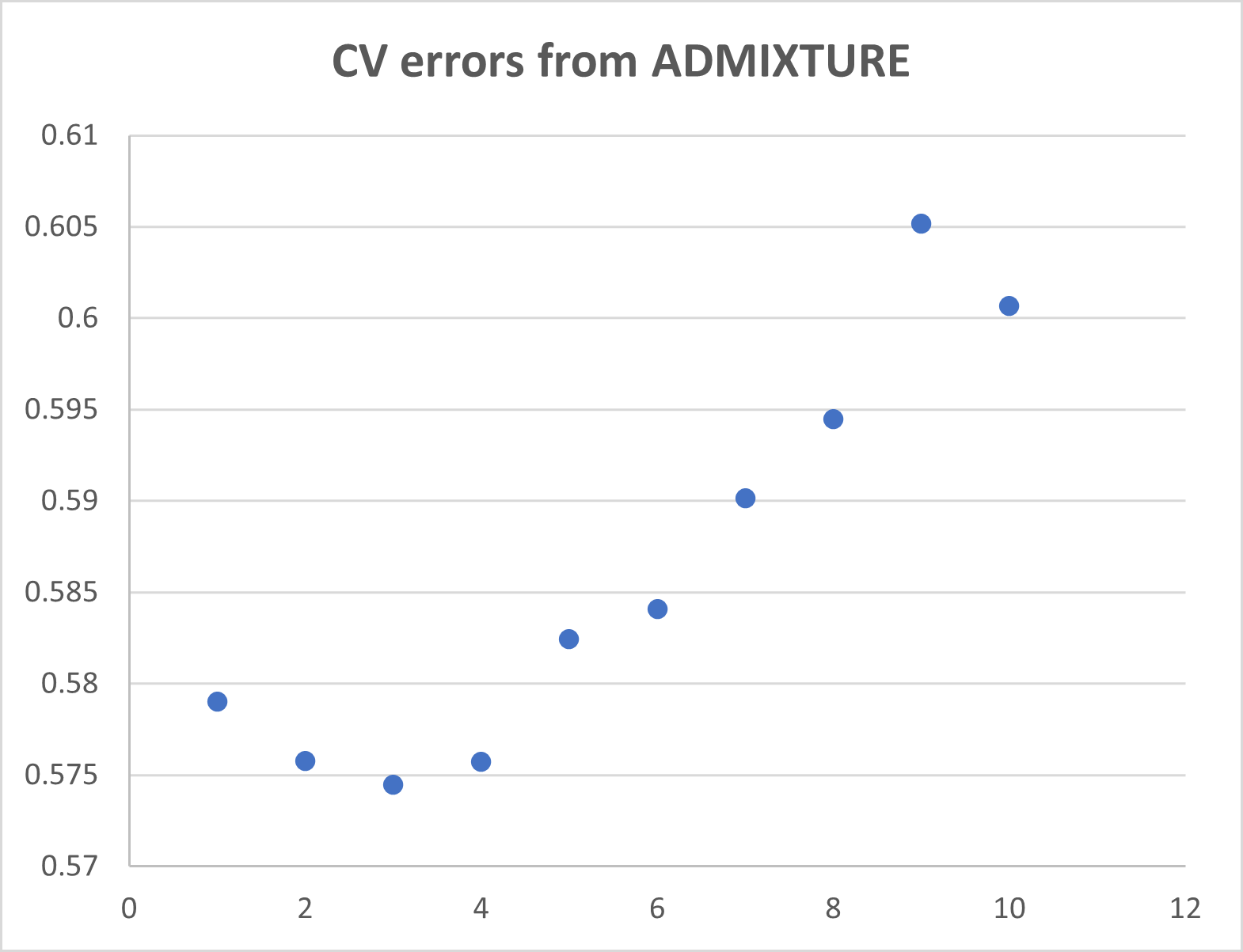

**Figure S3.** Cross-validation (CV) error estimates from ADMIXTURE, used to determine the optimal number of genetic clusters in the Tornio-Kalix River system (n = 361 samples, n = 20,745 SNPs).

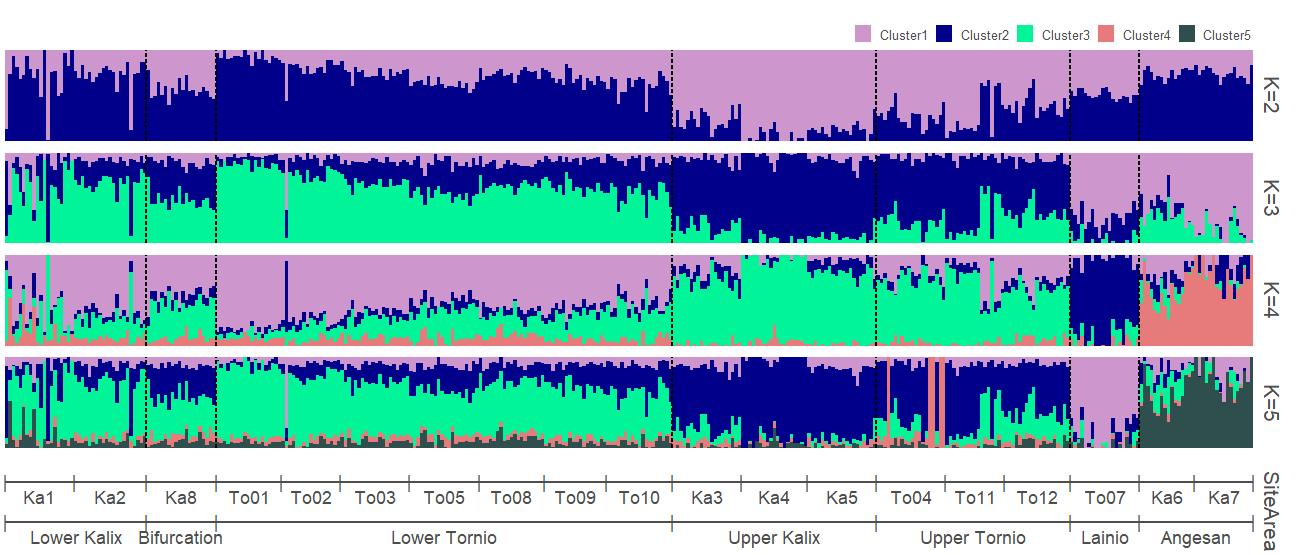

**Figure S4a.** ADMIXTURE coefficients for *K* = 2 to K = 5 (each vertical line depicts an individual) in the Tornio-Kalix salmon stock (n = 361 samples, 20,745 SNPs).

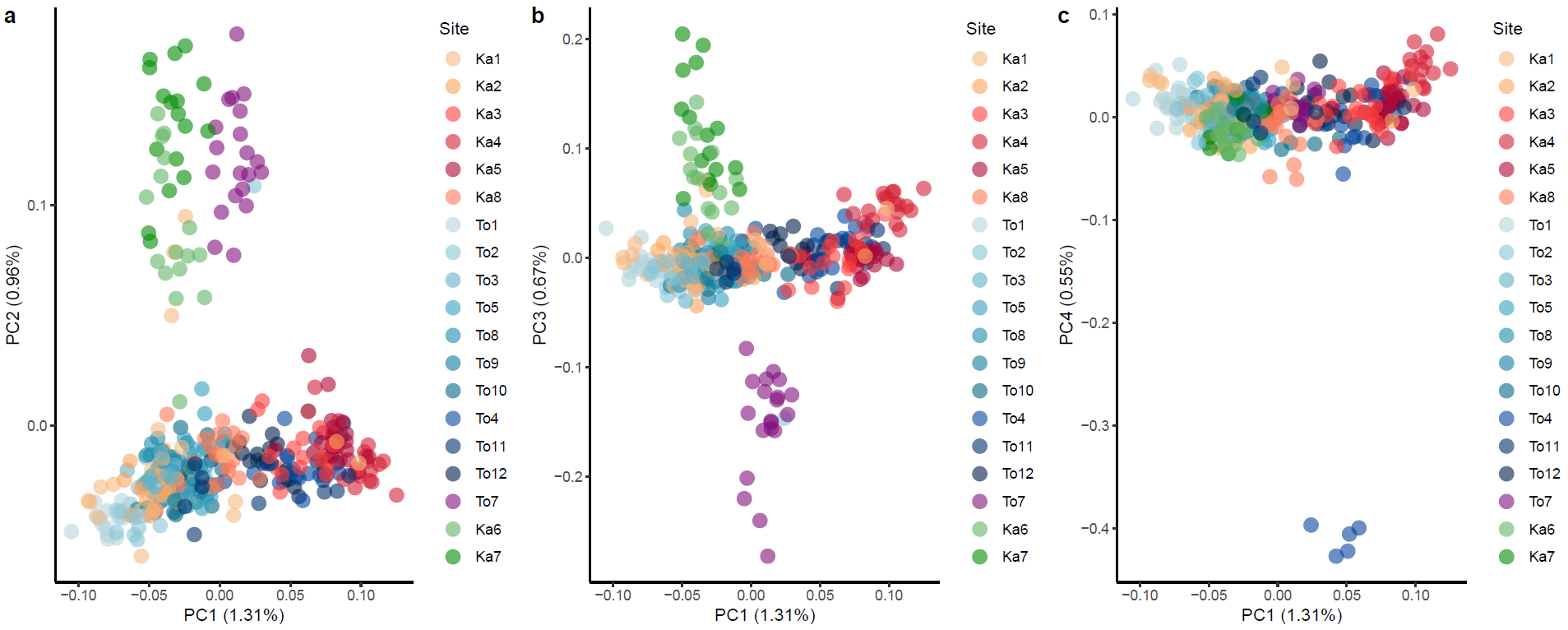

**Figure S4b.** Principal component analysis (PCA) visualising the genetic population structure of the Tornio-Kalix salmon stock (n = 361 samples, 20,745 SNPs), showing **a** PC1 and PC2, **b** PC1 and PC3, and **c** PC1 and PC4. The proportion of variance explained by each principal component is indicated on the axis labels. Each point represents an individual, and their distribution on the axes depicts their genetic distance from each other.

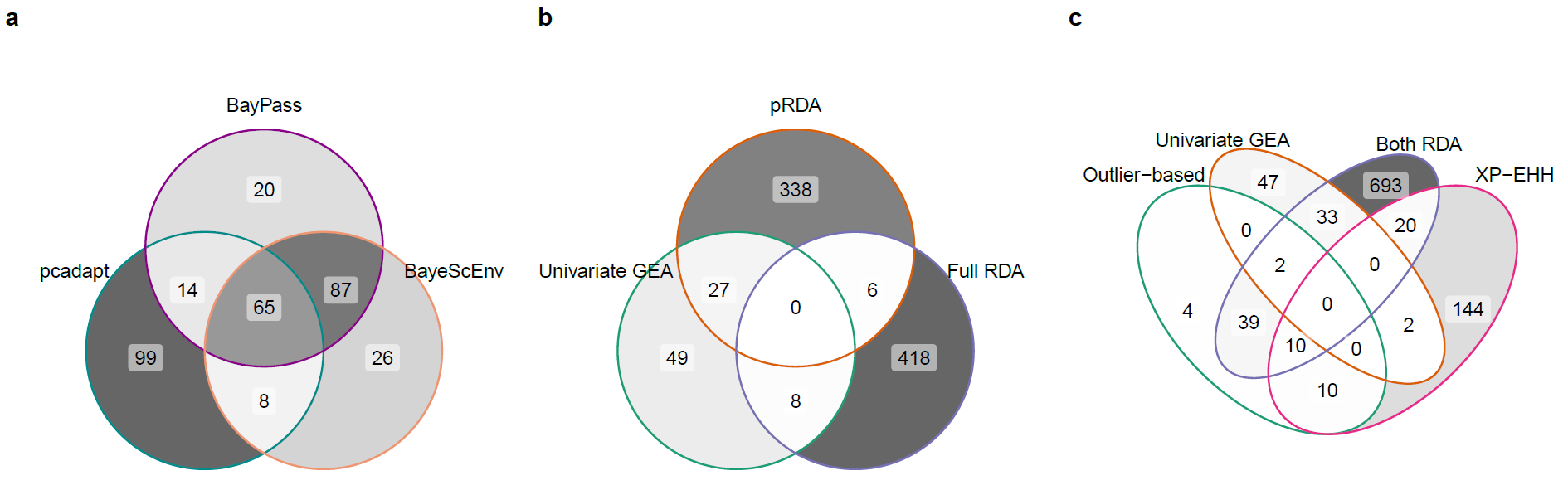
**Figure S5.** Venn diagrams of overlaps among top-ranked/candidate SNPs identified by the different types of analyses. Overlapping **a** top-ranked SNPs from the three outlier-based analyses (i.e. the 186 top-ranked SNPs from each test), and candidate SNPs **b** from the GEA analyses, and **c** from all analyses (i.e. outlier-based, univariate and multivariate GEA, and haplotype-based analyses).

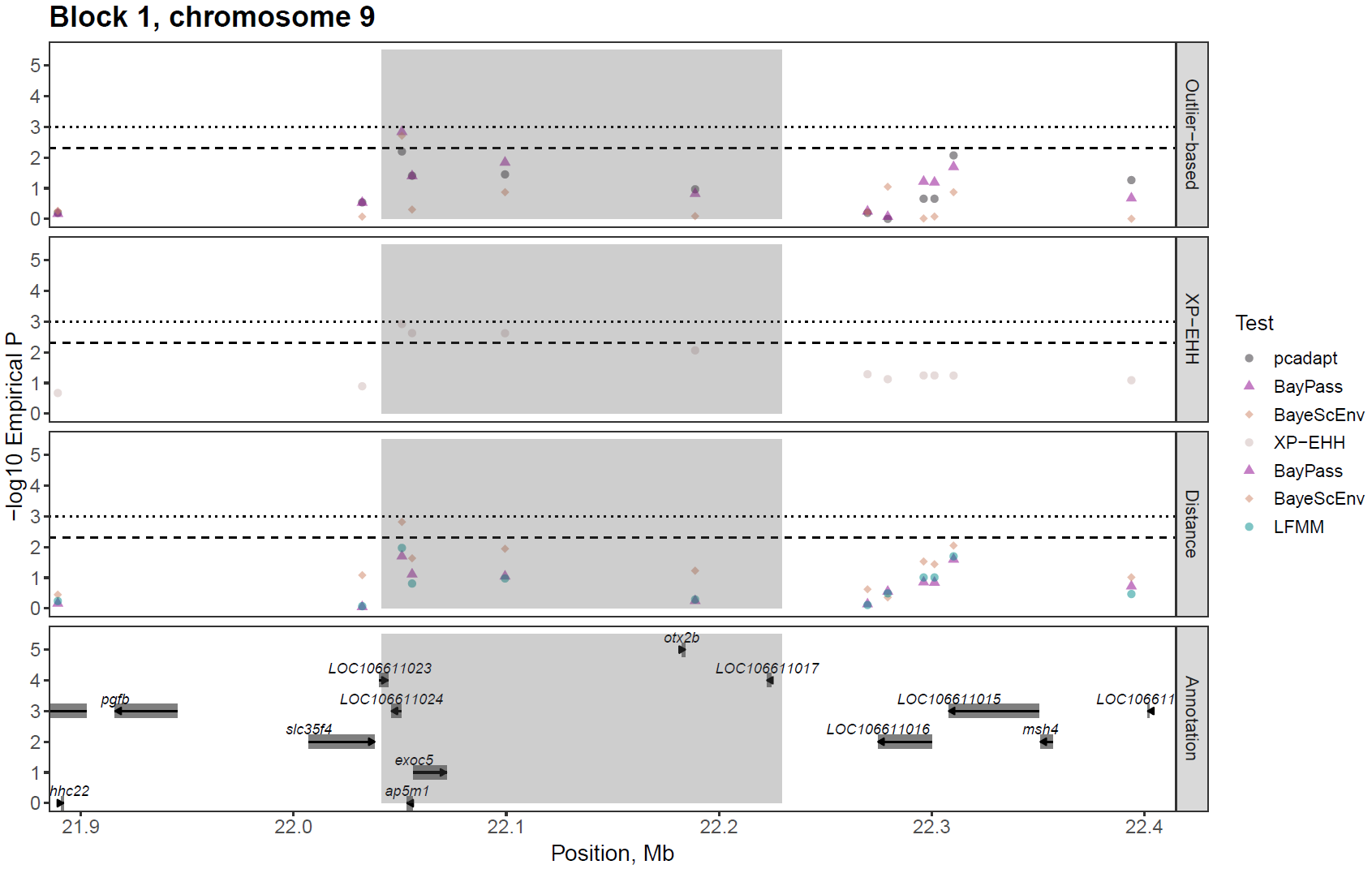

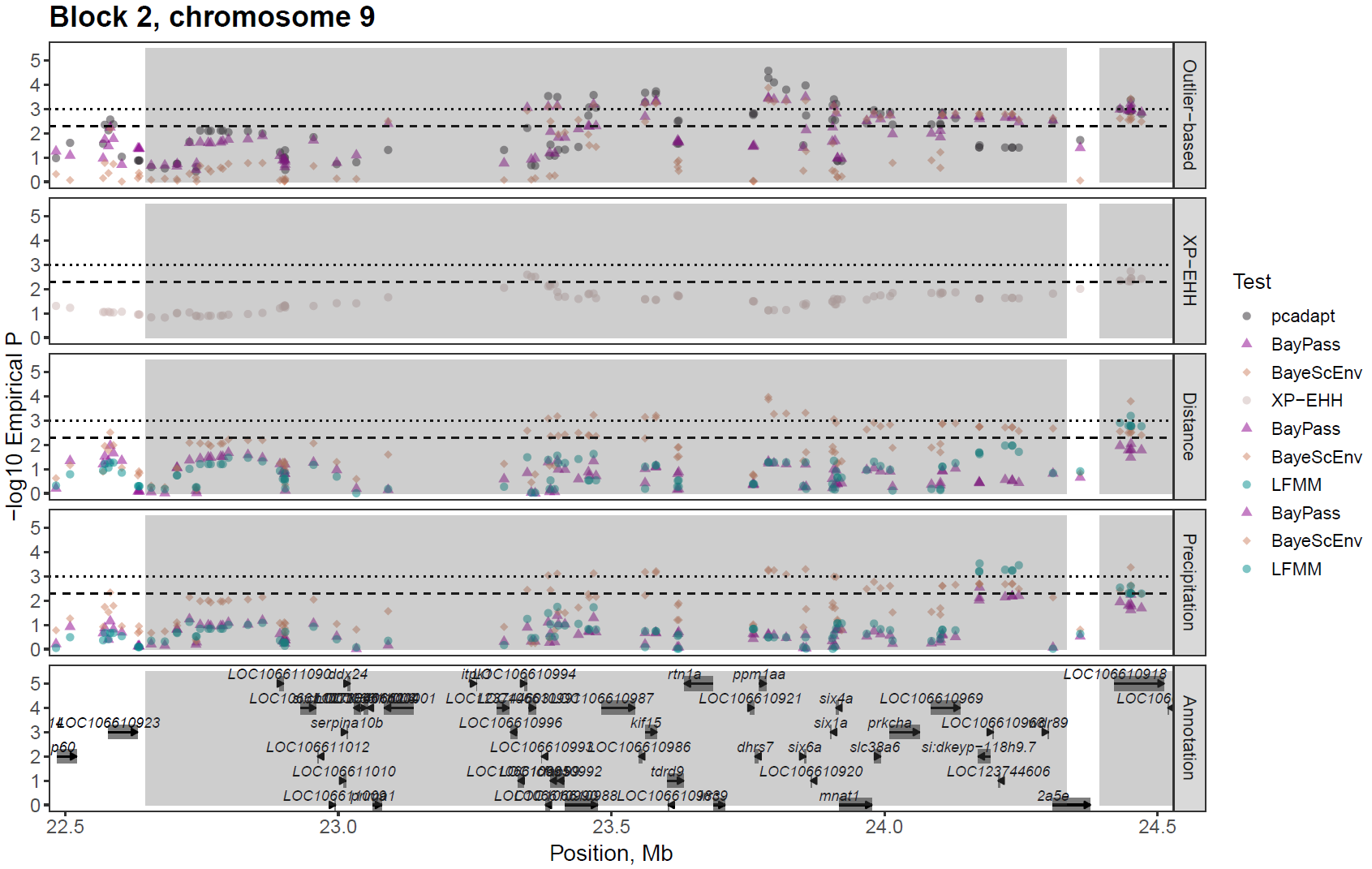

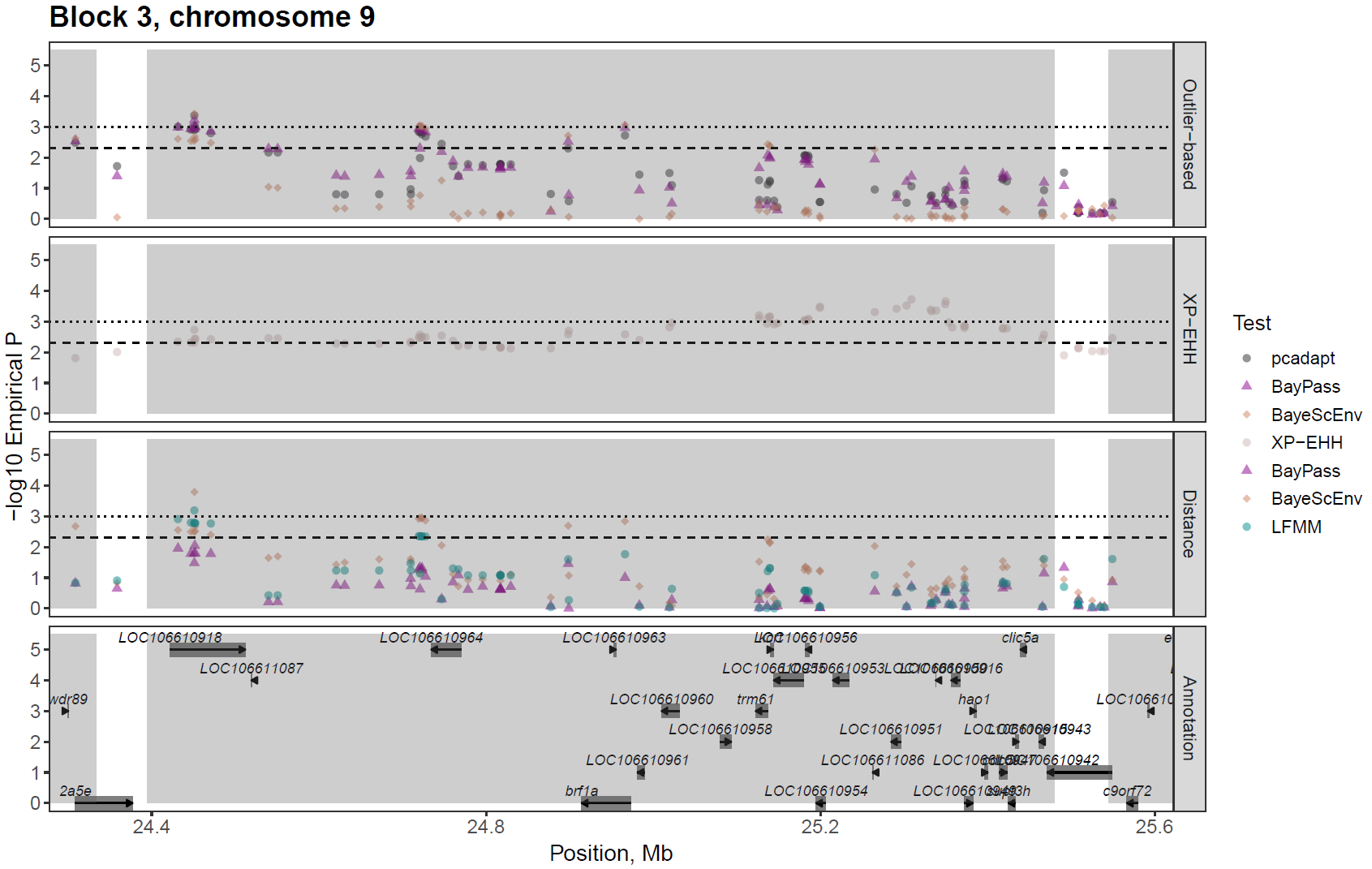

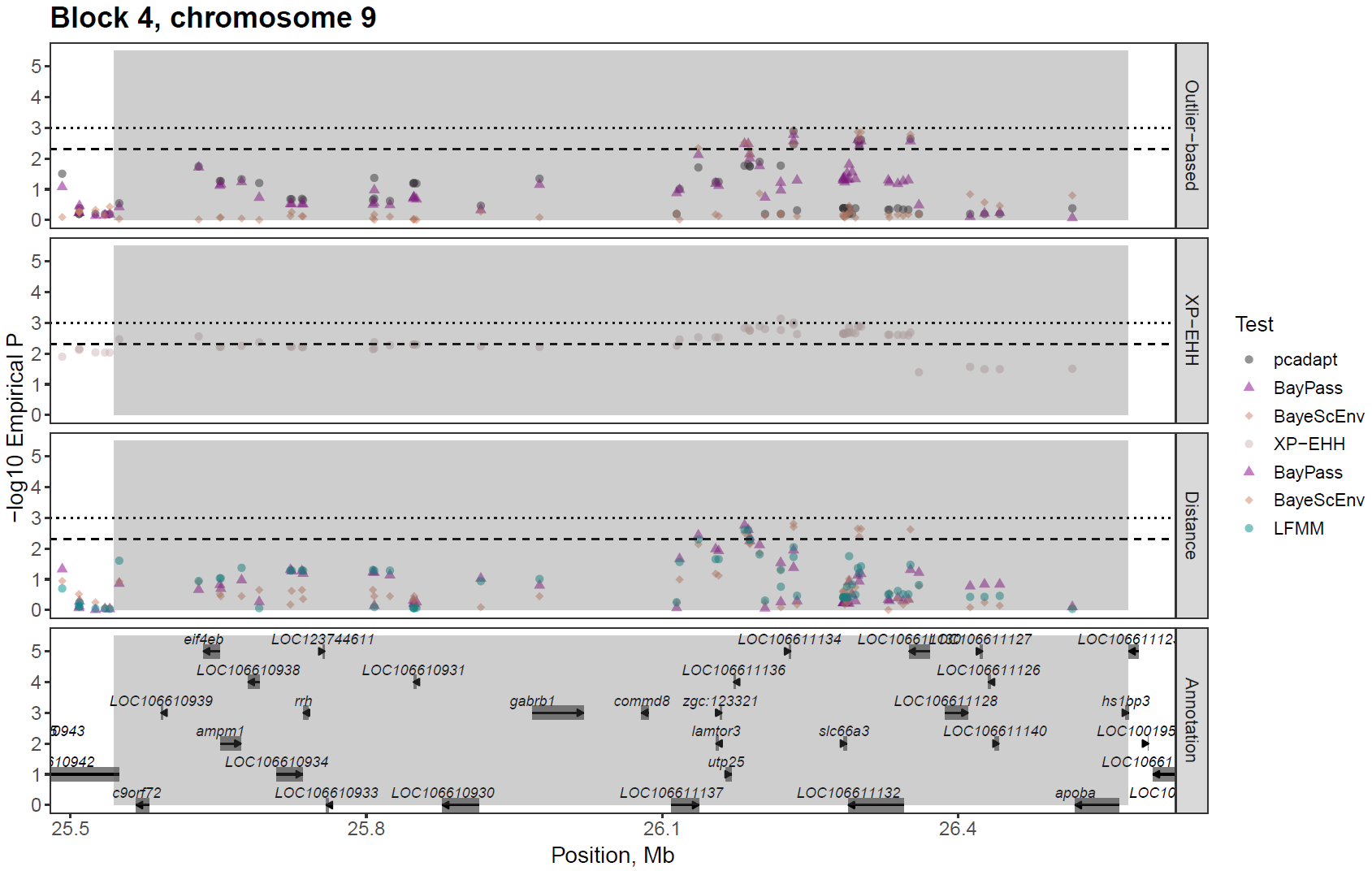

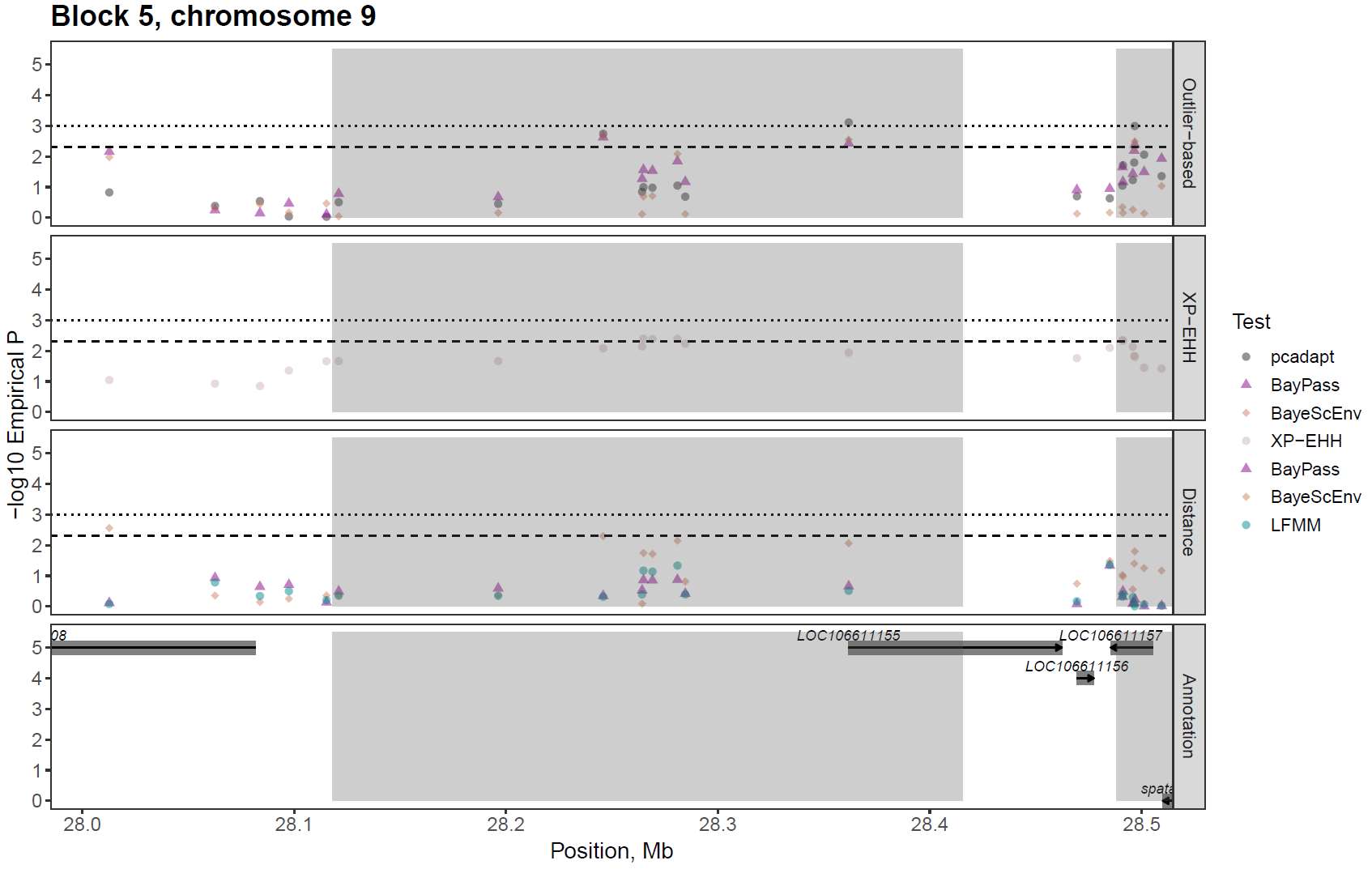

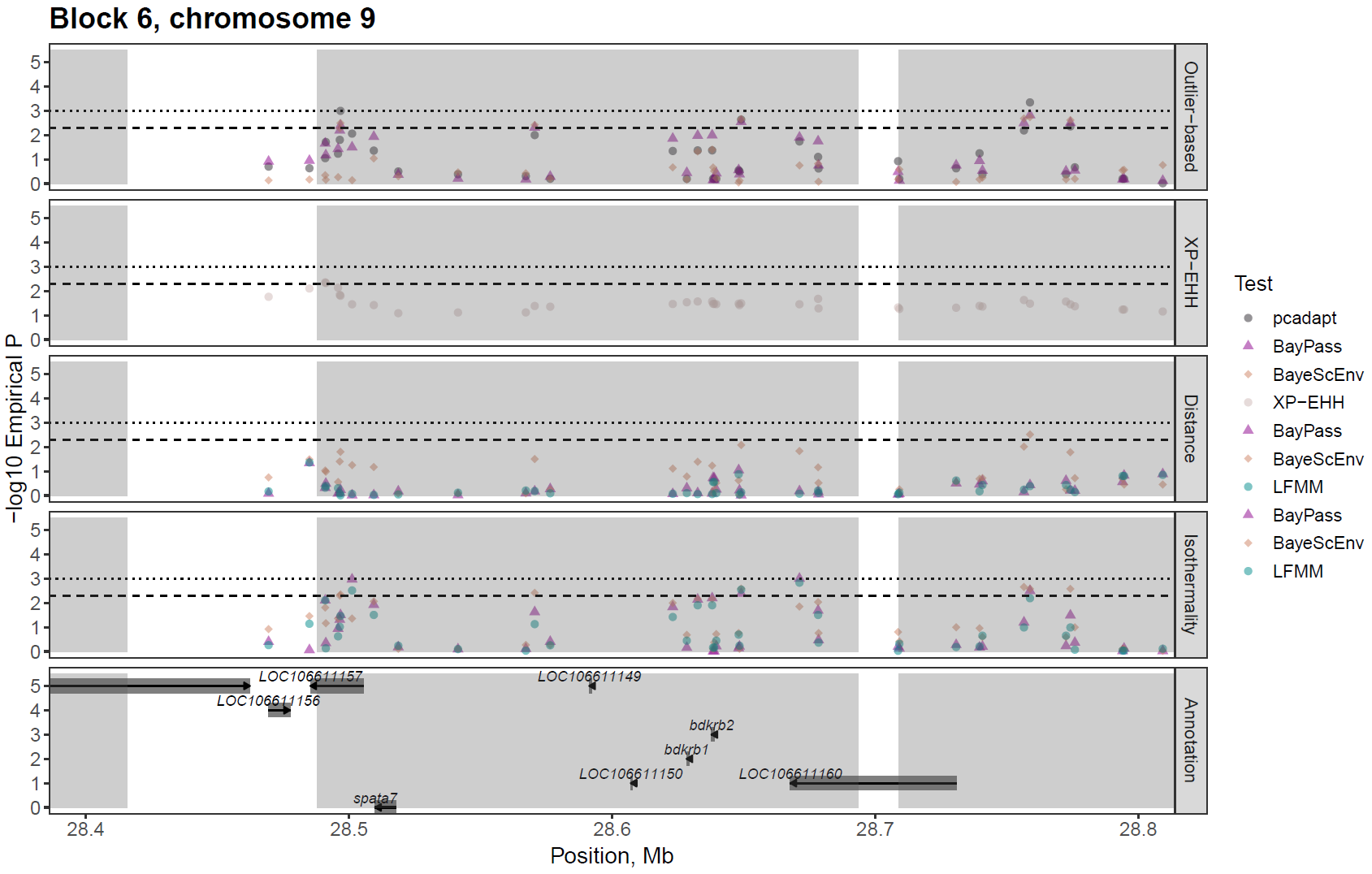

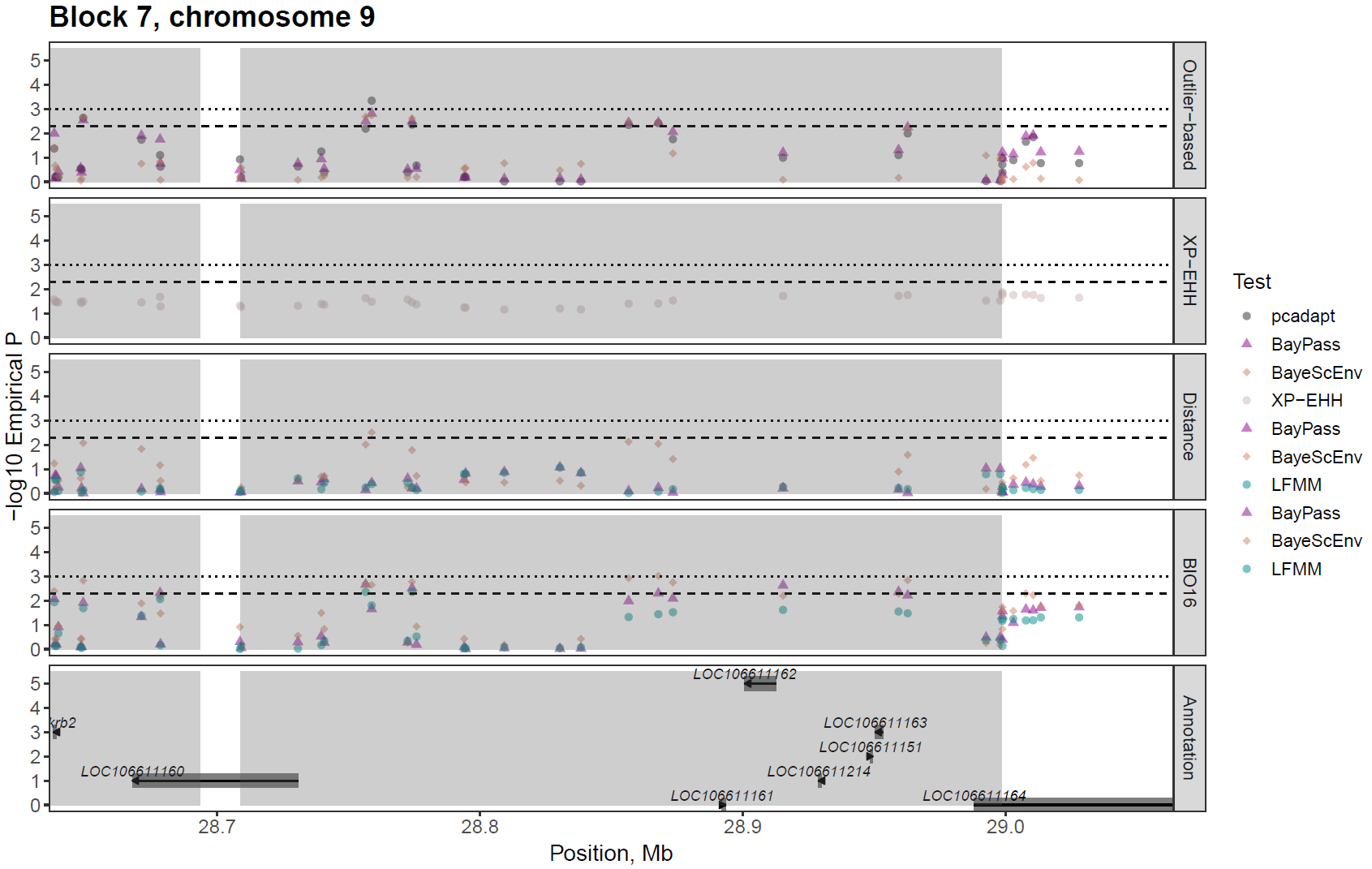

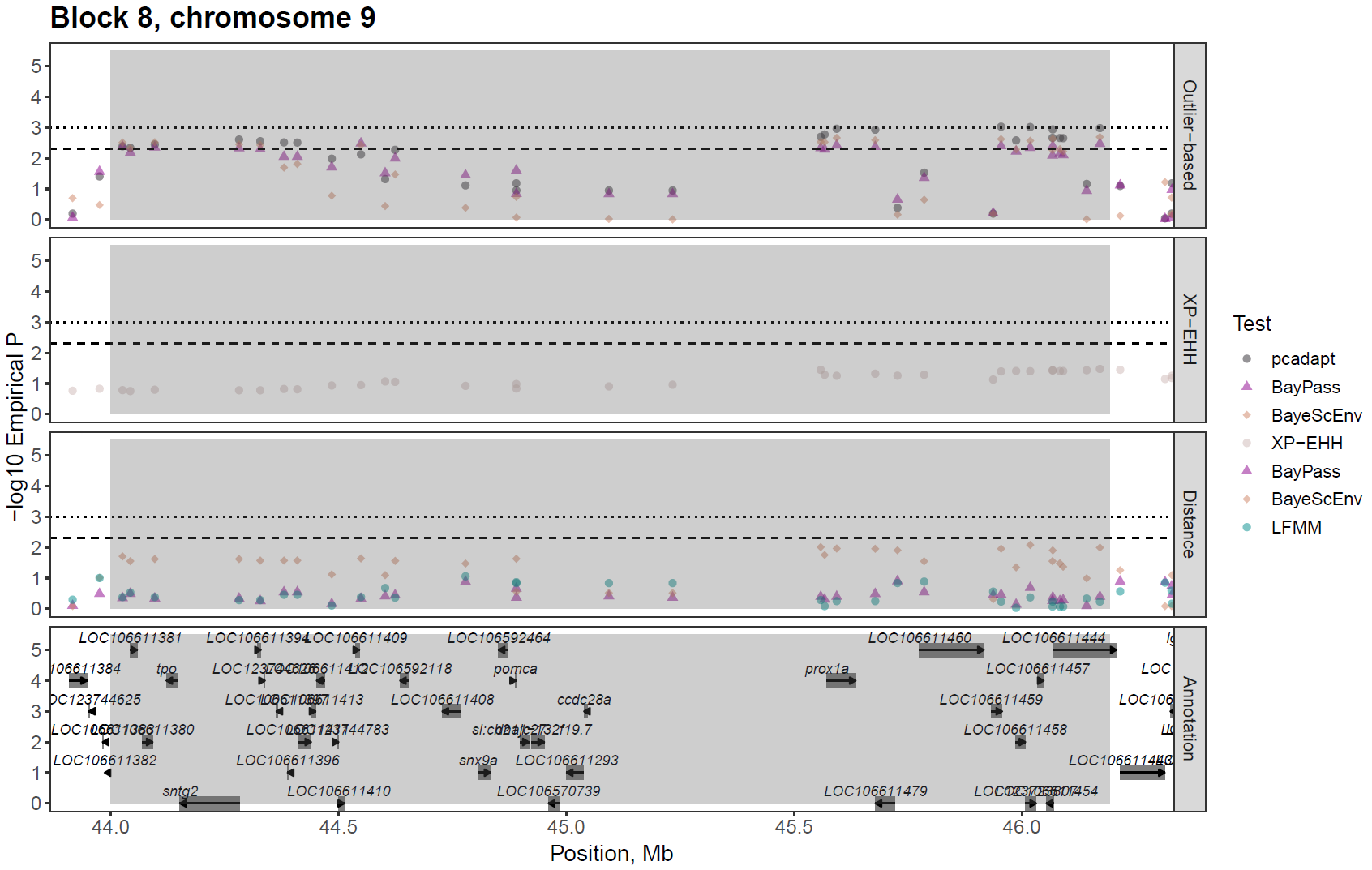

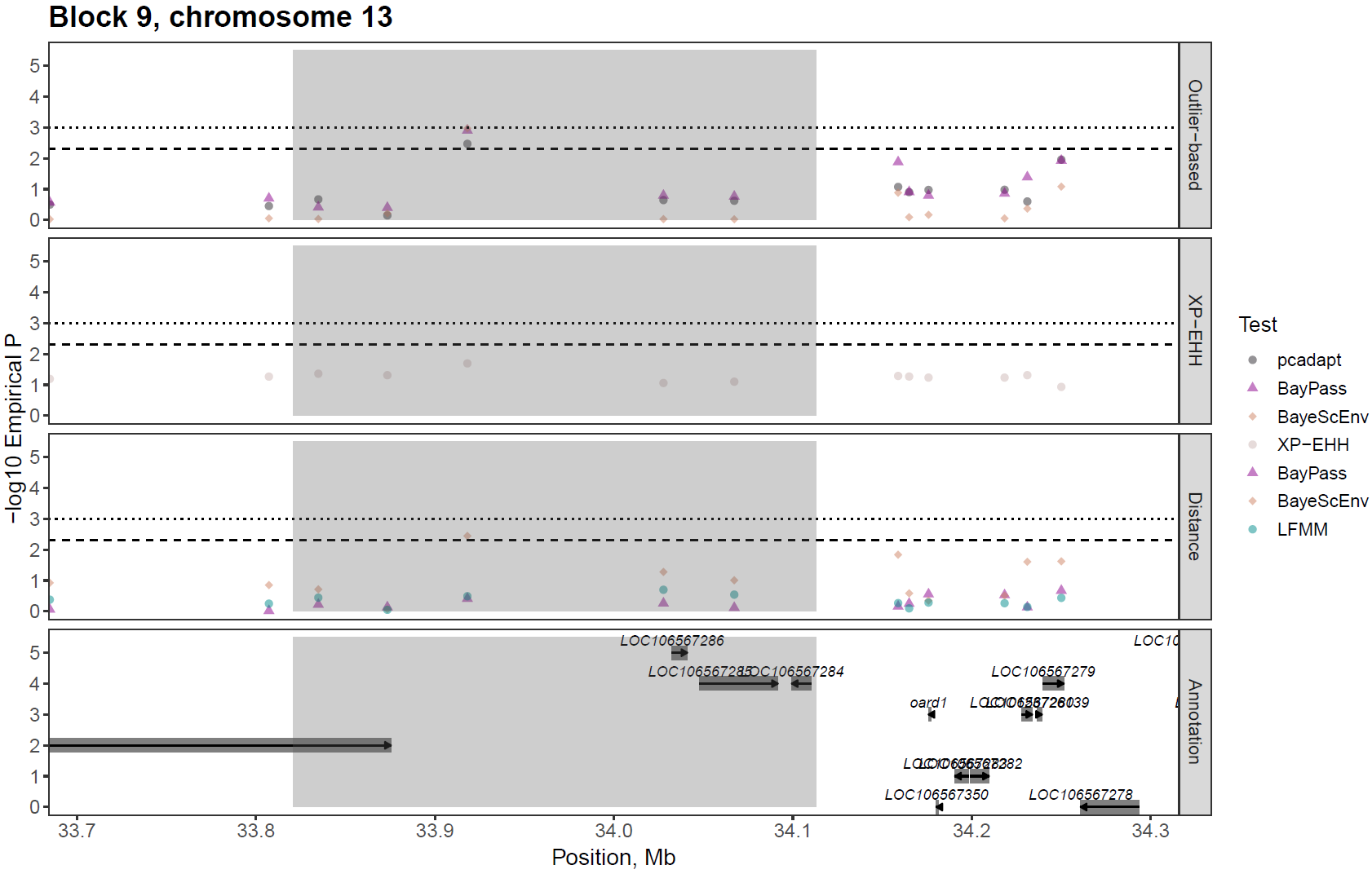

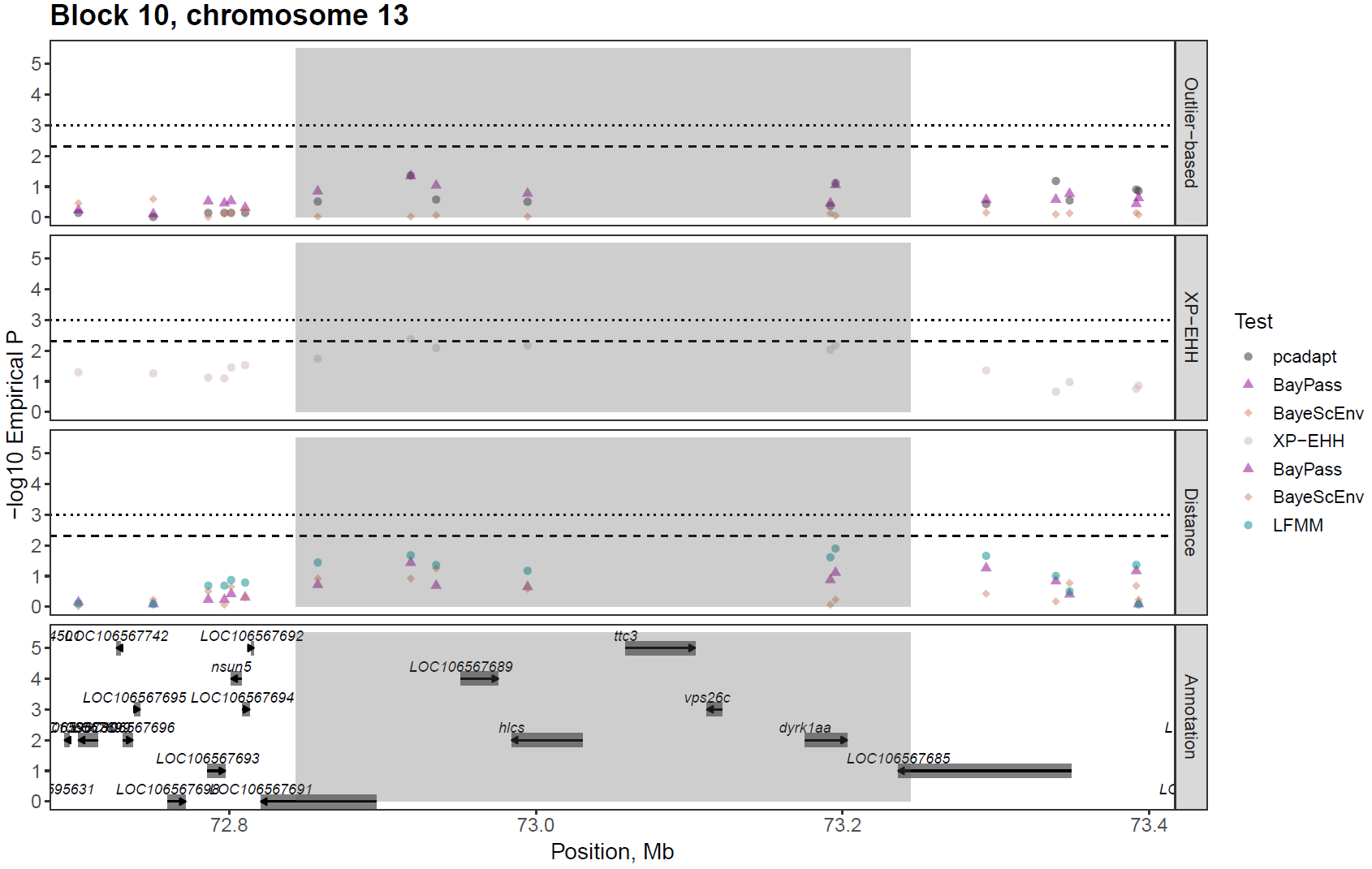

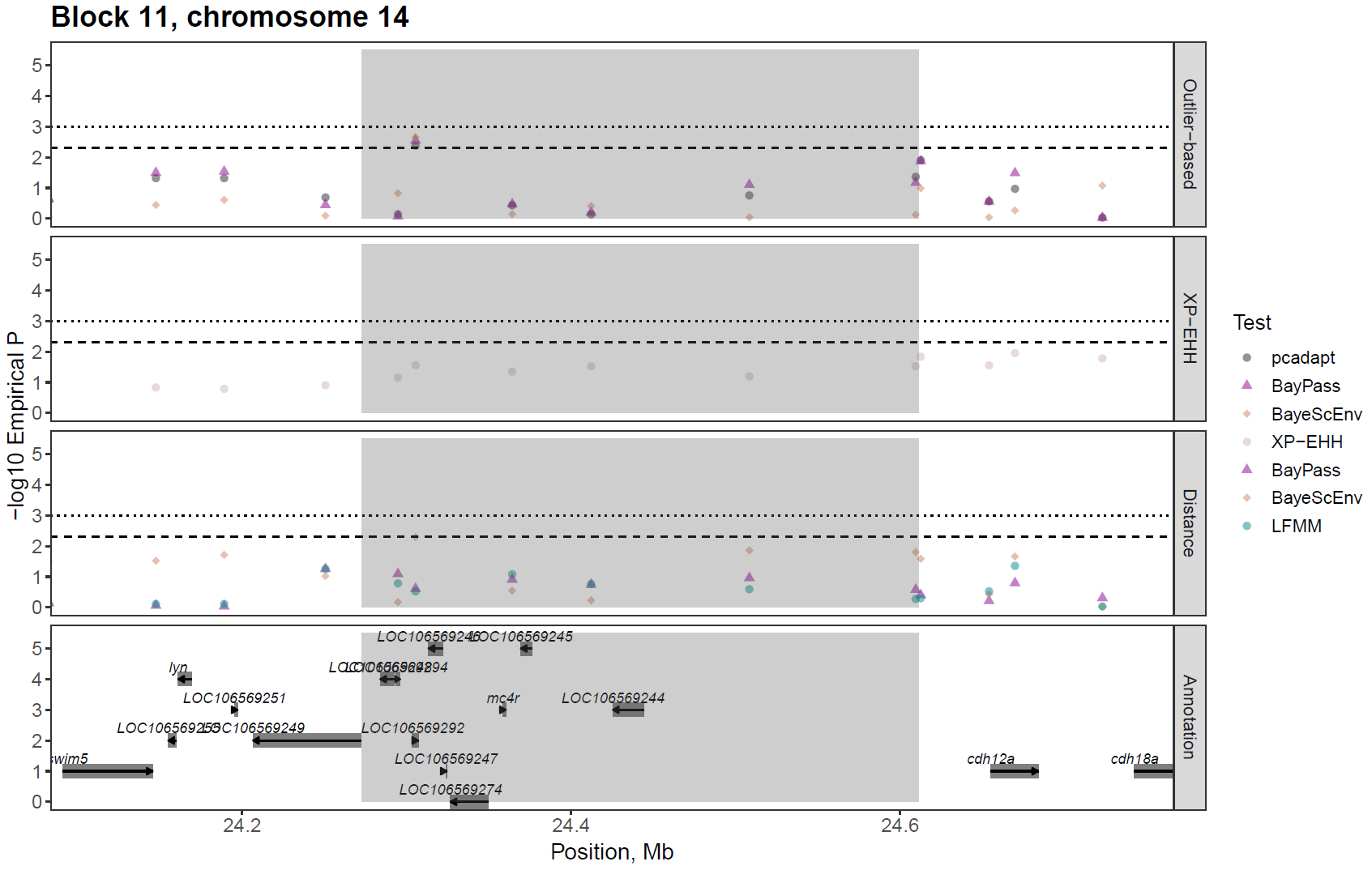

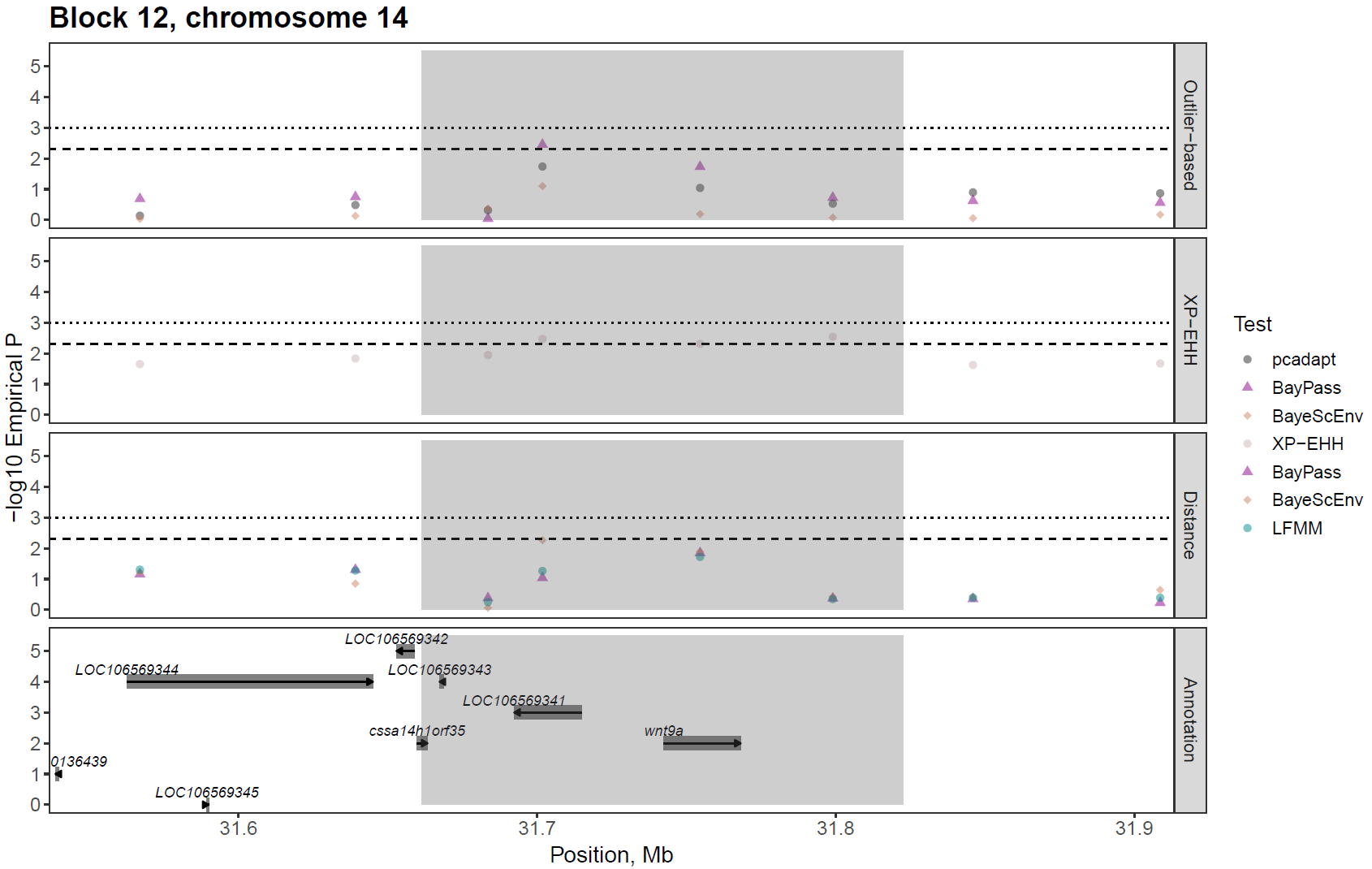

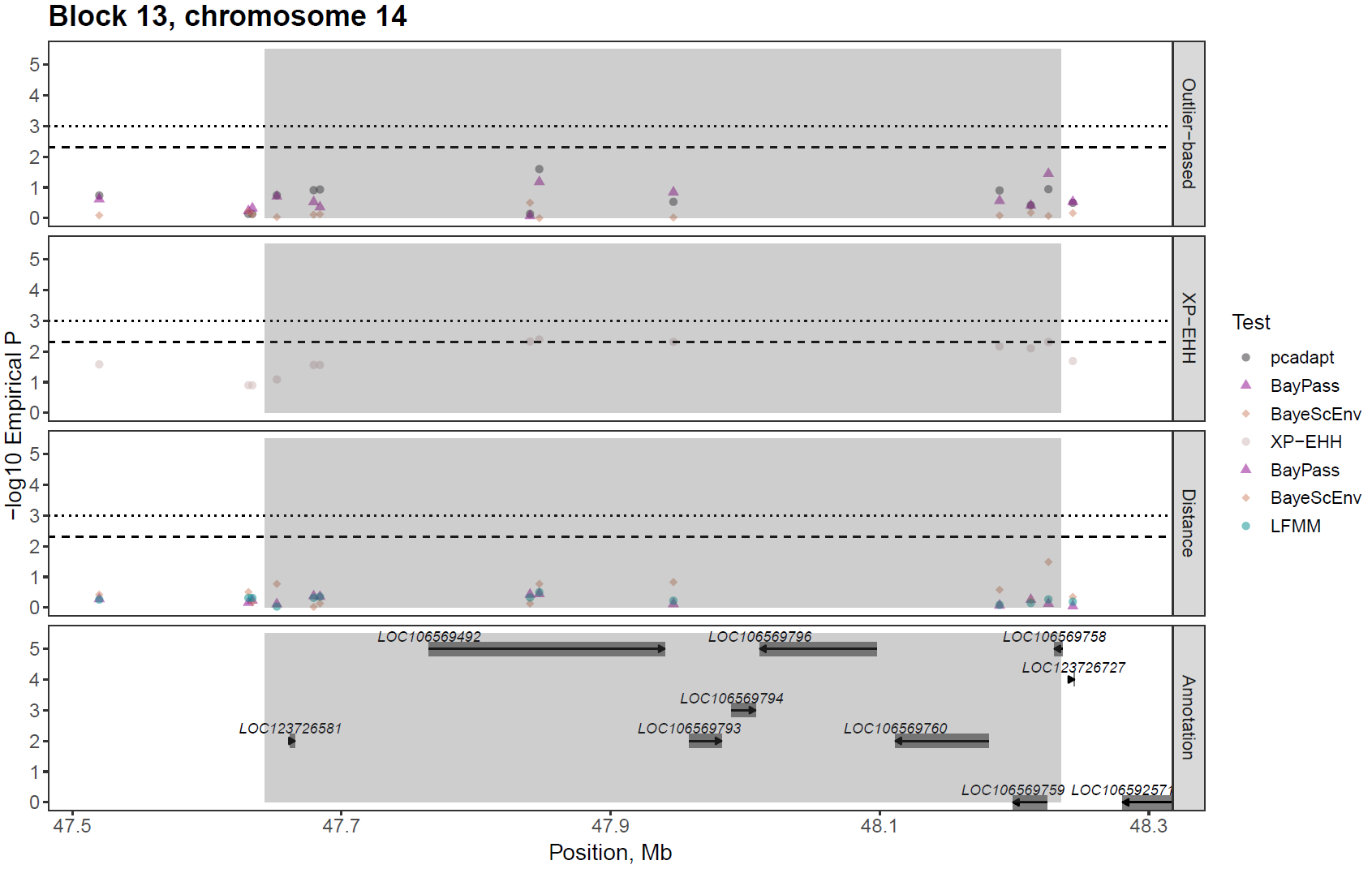

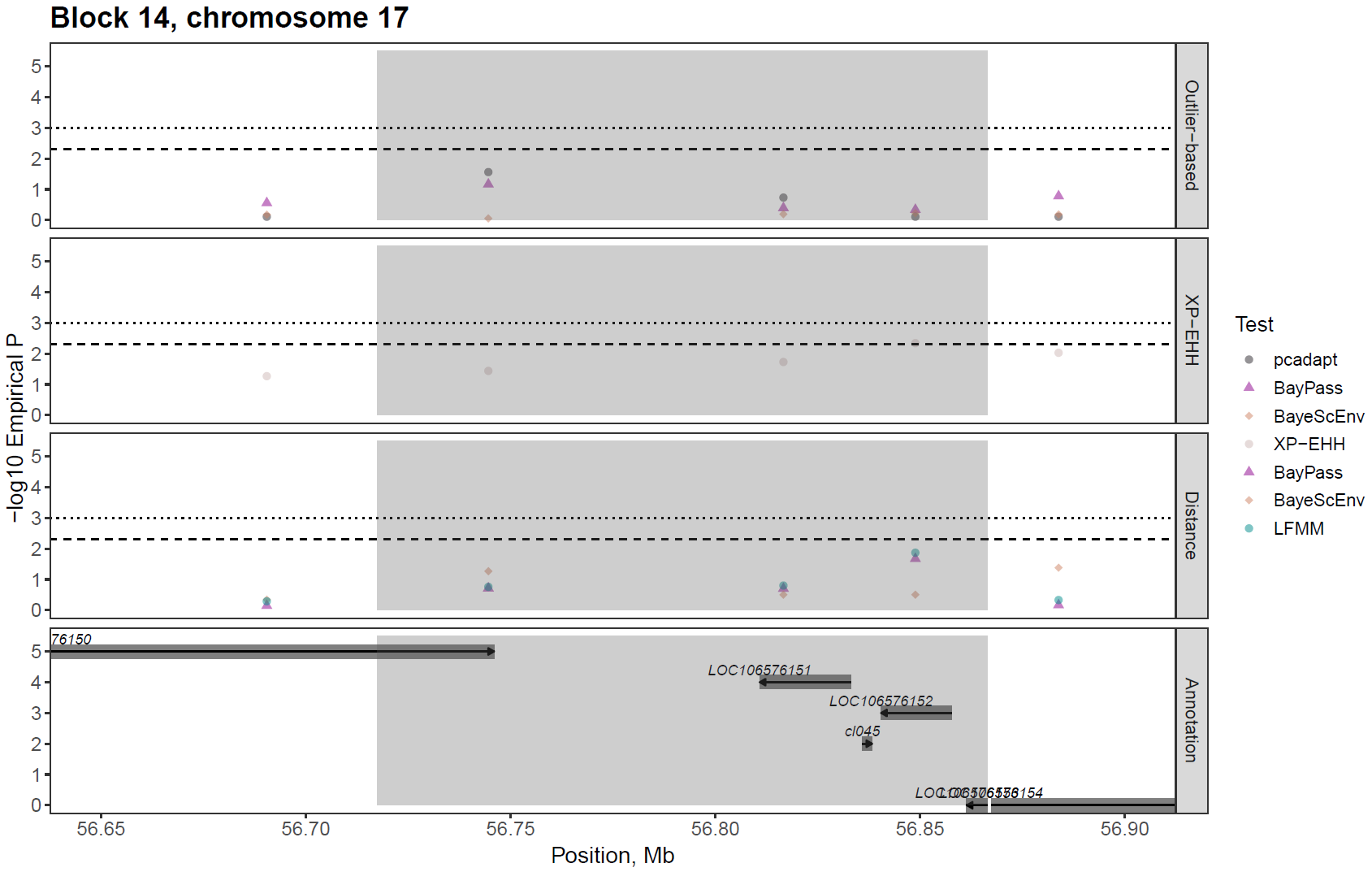

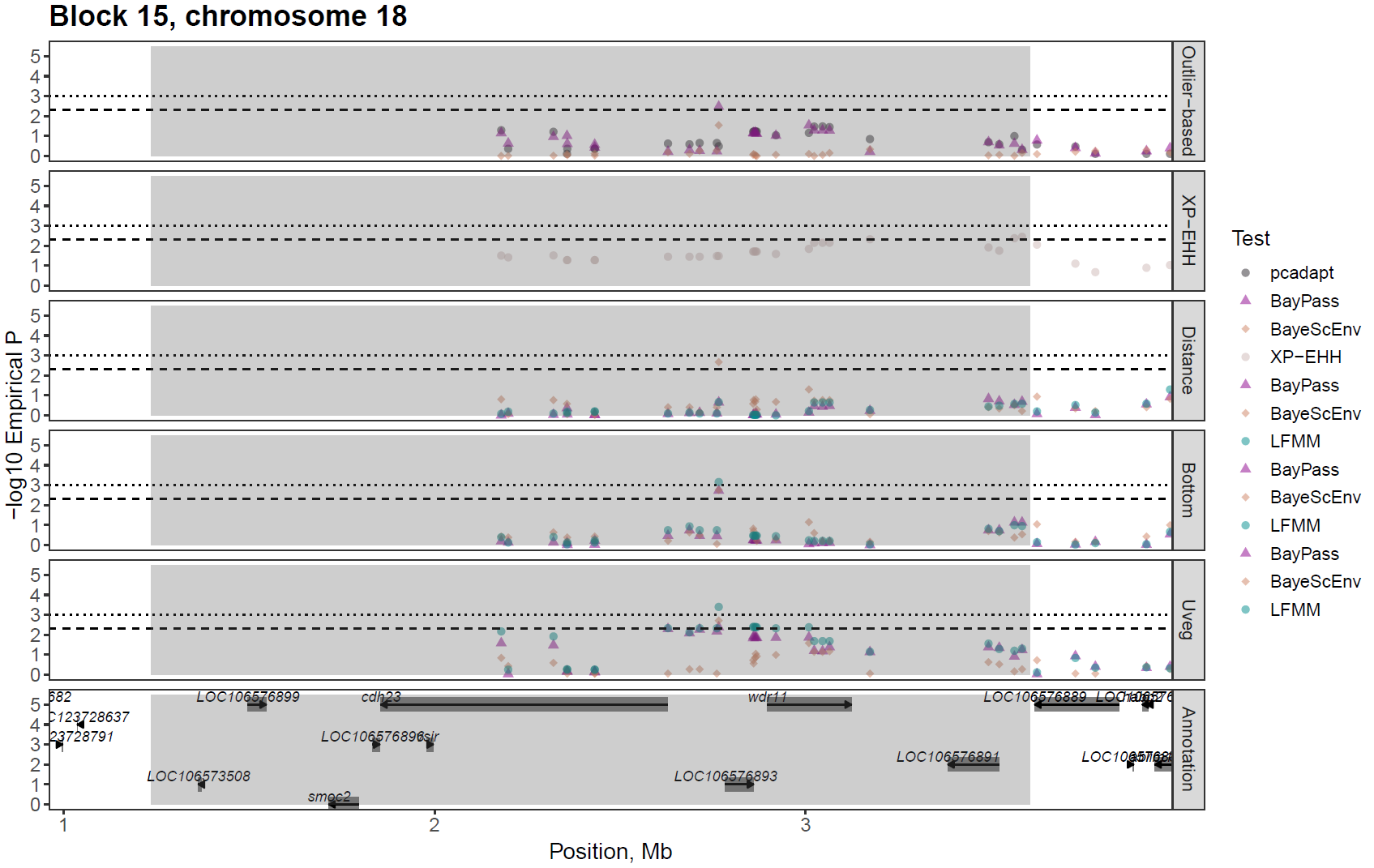

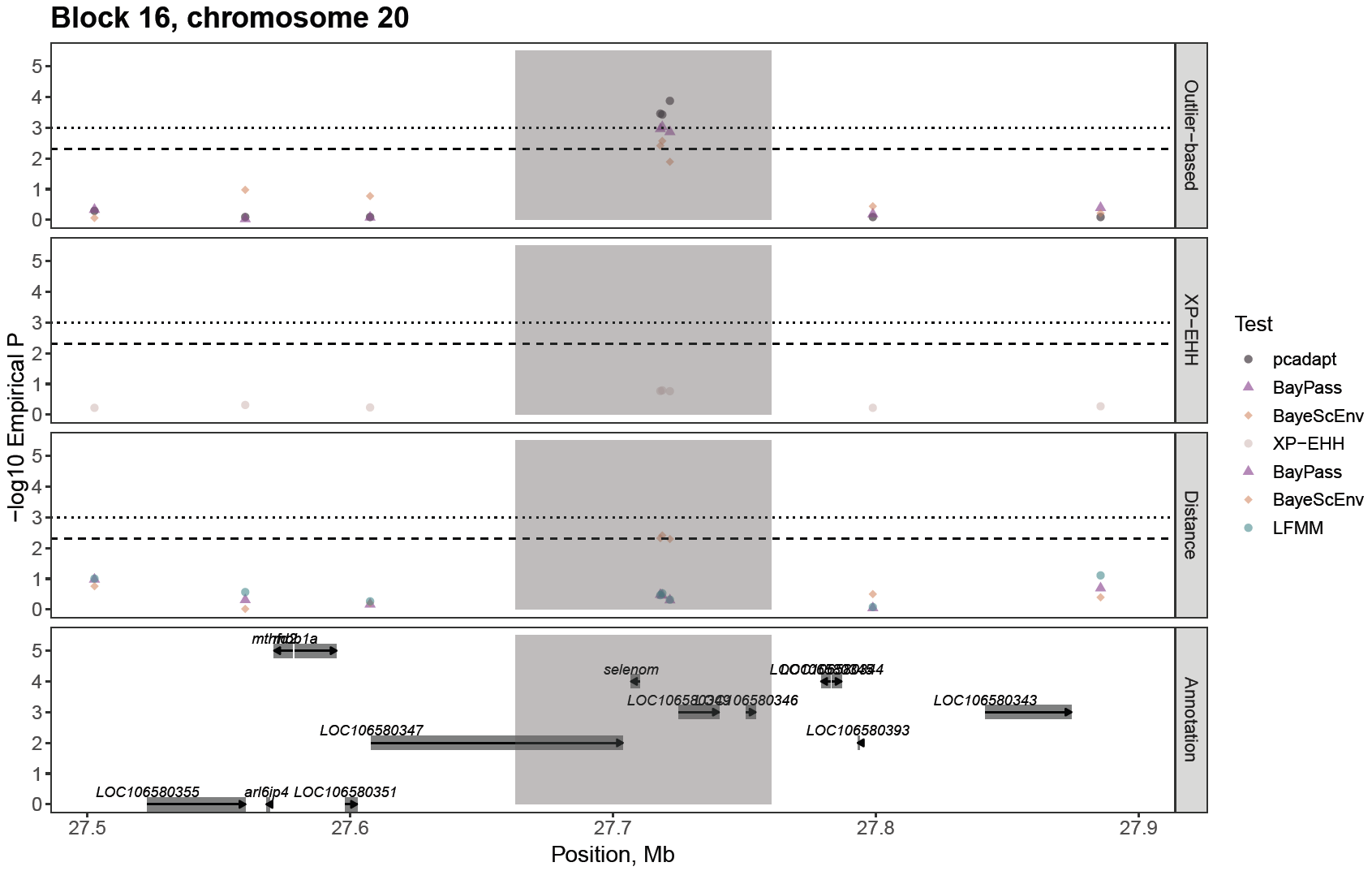

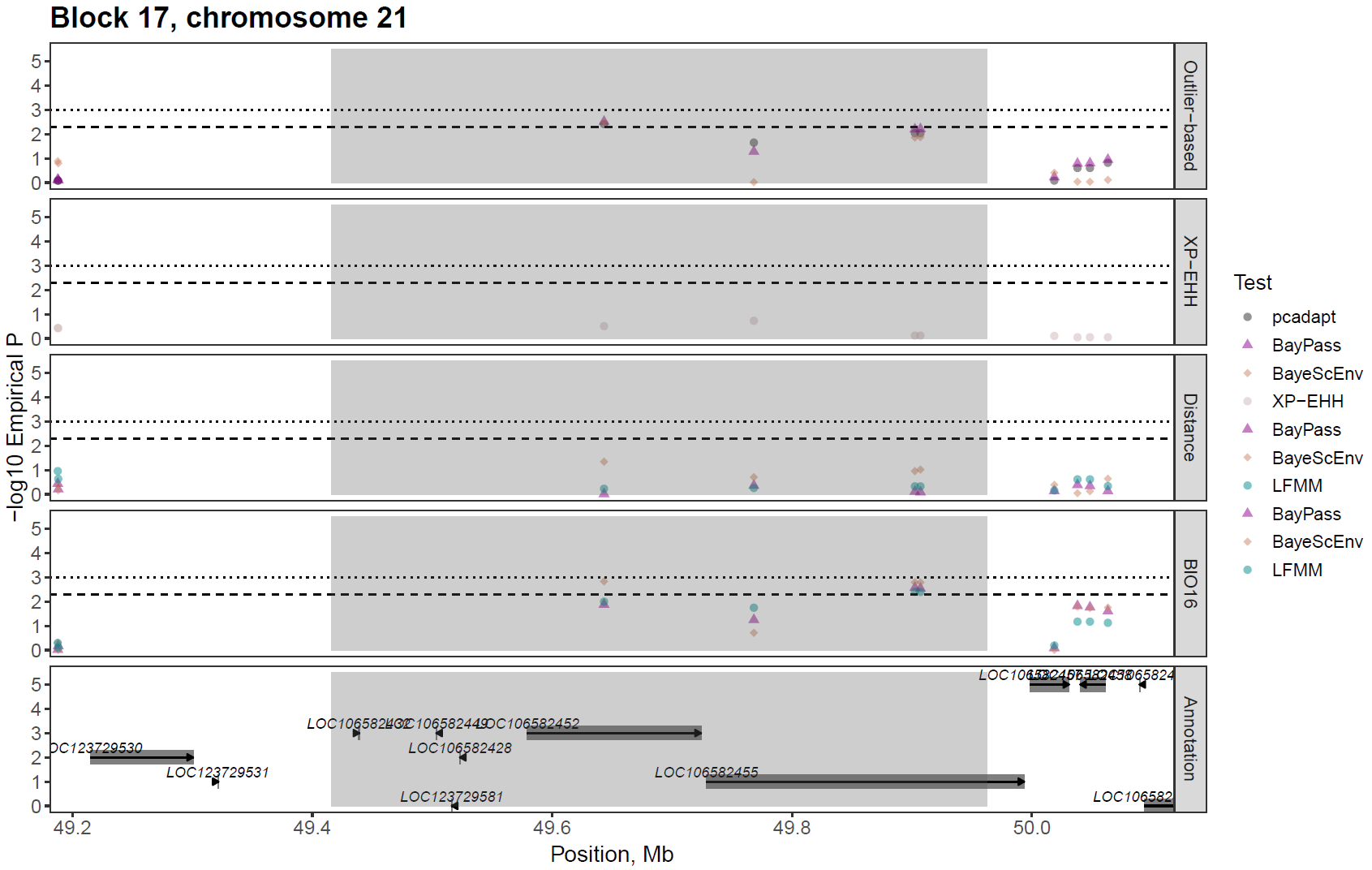

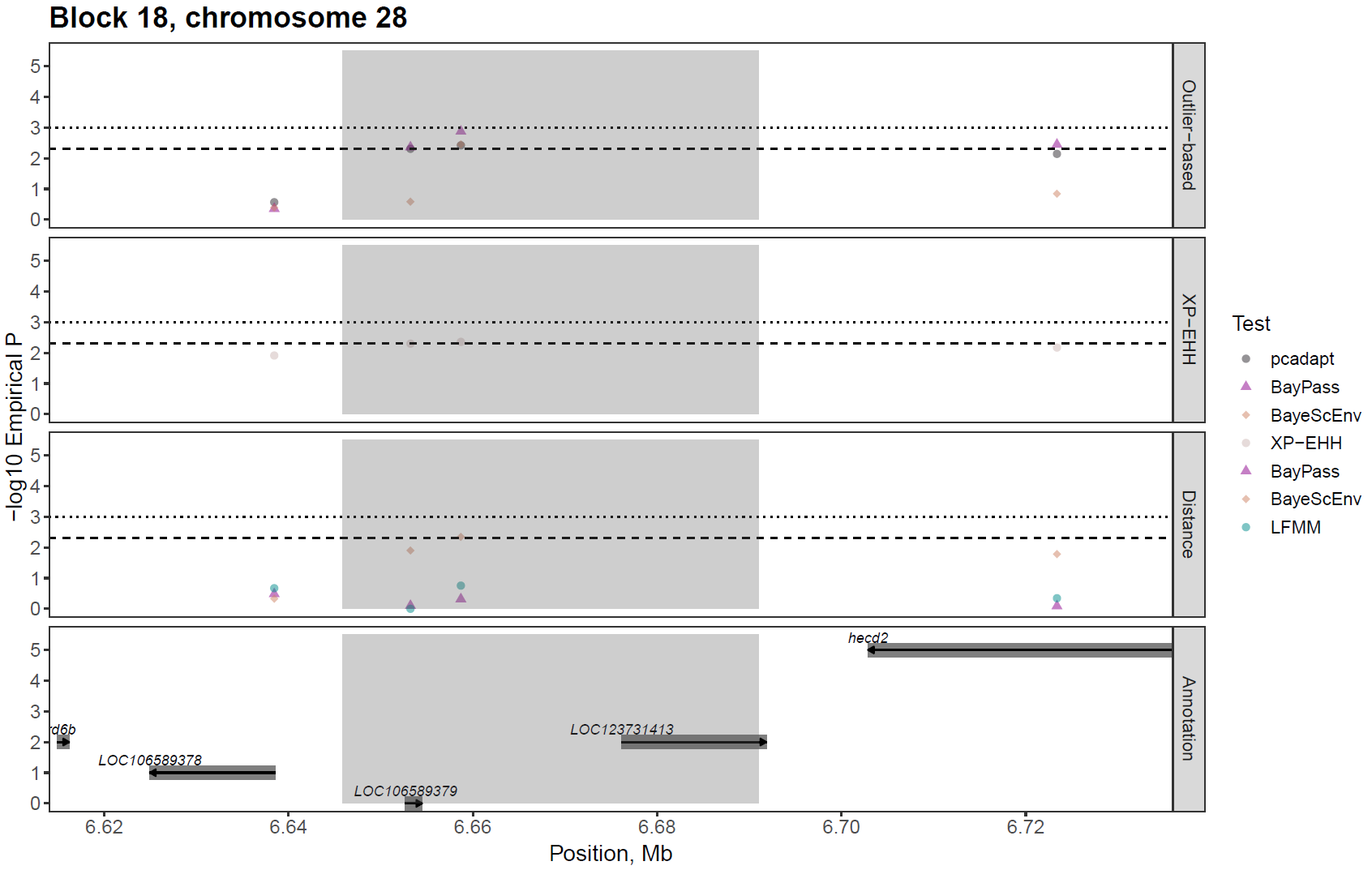

**Figures S6a-r (above).** Signatures of local selection on the 18 candidate haploblocks identified in this study (Table 3). The dashed and dotted lines indicate empirical p < 0.005 and p < 0.001, respectively (empirical p = SNP rank/total number of tests). The grey rectangles show the boundaries of candidate haploblocks. The panels from top to bottom show empirical p-values for outlier-based tests, XP-EHH test, univariate GEA analyses of distance from river mouth (also a proxy for e.g. temperature-related variation) and of the environmental variable(s) associated with candidate SNPs in the haploblock in both univariate and multivariate GEA analyses.

**References (Table S5)**
